# Structure of the Vesicular Stomatitis Virus L Protein in Complex with Its Phosphoprotein Cofactor

**DOI:** 10.1101/792960

**Authors:** Simon Jenni, Louis-Marie Bloyet, Ruben Diaz-Avalos, Bo Liang, Sean P. J. Whelan, Nikolaus Grigorieff, Stephen C. Harrison

## Abstract

The large (L) proteins of non-segmented, negative-strand RNA viruses are multifunctional enzymes that produce capped, methylated and polyadenylated mRNAs and replicate the viral genome. A phosphoprotein (P), required for efficient RNA-dependent RNA polymerization from the viral ribonucleoprotein (RNP) template, regulates function and conformation of the L protein. We report the structure of vesicular stomatitis virus L in complex with its P cofactor determined by electron cryomicroscopy at 3.0 Å resolution, enabling us to visualize bound segments of P. The contacts of three P segments with multiple L domains show how P induces a closed, compact, initiation-competent conformation. Binding of P to L positions its N-terminal domain adjacent to a putative RNA exit channel for efficient encapsidation of newly synthesized genomes with the nucleoprotein and orients its C-terminal domain to interact with the RNP template. The model shows that a conserved tryptophan in the priming loop can support the initiating 5’-nucleotide.

## INTRODUCTION

The large (L) protein encoded by the genomes of nonsegmented, negative-sense (NNS) RNA viruses carries out all the various catalytic steps associated with transcription and replication. A virally encoded phosphoprotein (P) is an essential cofactor, both for incorporation of L into virions and for regulating replication and transcription. In addition to its RNA-dependent RNA polymerase (RdRp) activity, L caps and methylates the 5’ ends of transcripts.

The vesicular stomatitis virus (VSV) genome encodes an untranslated (and uncapped) 5’ leader sequence and five proteins, in the order of N, P, M, G, and L. The template for transcription is a full-length ribonucleoprotein (RNP) – that is, a genome-sense RNA fully coated with protein N. Each N subunit accommodates nine nucleotides of RNA in a groove along the waist of an elongated, two-lobe protein. Transcription of successive genes (with about 70% efficiency) initiates upon termination and polyadenylation of the upstream transcript, produced by stuttering on a U7 sequence at the end of each gene (Iverson and Rose, 1981).

We described four years ago the structure of a VSV L-P complex from a cryo-EM reconstruction at 3.8 Å resolution (Liang et al., 2015). The multifunctional L protein, a single, 2109 amino-acid long polypeptide chain, folds into three catalytic and two structural domains (Figure 1A and Data S1). The N-terminal RNA-dependent RNA polymerase domain (RdRp) together with a capping domain (Cap), which follows in primary sequence and has polyribonucleotidyl transferase (PRNTase) activity, form the core of the structure (Figure 1B). Negative-stain electron microscopy (EM) images had shown previously that in the absence of P, the three remaining domains – the connector domain (CD), the methyltransferase (MT), and the C-terminal domain (CTD) – have no fixed position with respect to the RdRp-Cap core structure, and the full molecular model from cryo-EM showed particularly long linker segments between Cap and CD and between CD and MT (Liang et al., 2015; Rahmeh et al., 2010).

**Figure 1.**
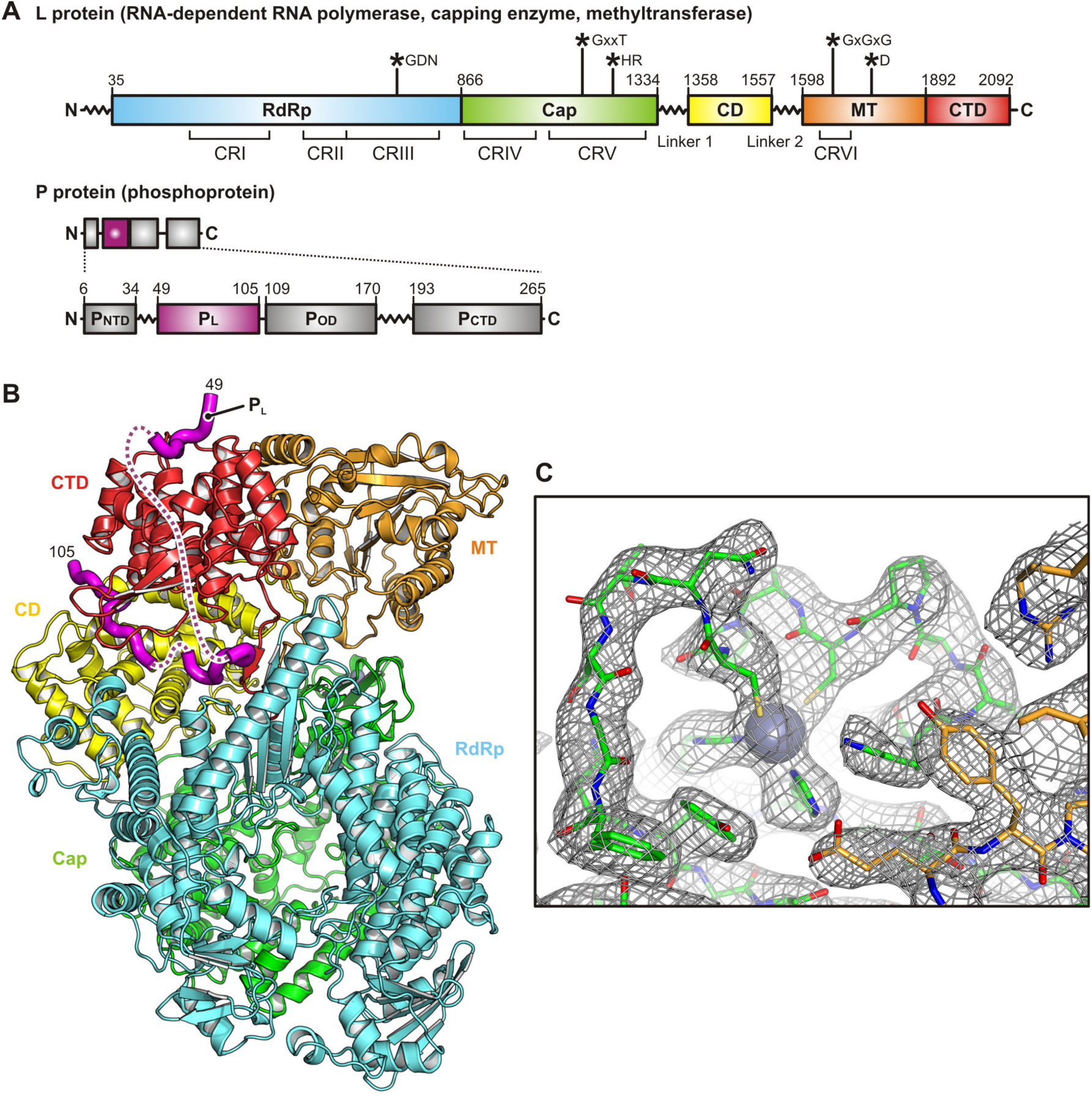
Structure of the VSV L Protein in Complex with the Phosphoprotein. (A) Linear domain organization scheme for the VSV L and P proteins. Numbers for domain boundaries are shown. L protein domains are colored as follows: RNA-dependent RNA polymerase domain (RdRp), cyan; capping domain (Cap), green; connector domain (CD), yellow; methyltransferase (MT), orange; C-terminal domain (CTD), red. Catalytic residues are shown on top. CR I–VI are conserved regions within L proteins of non-segmented negative-strand (NNS) RNA viruses. The L protein scheme is adapted from (Liang et al., 2015). The P protein’s N-terminal domain (P_NTD_), oligomerization domain (P_OD_), and C-terminal domain (P_CDT_) are colored gray. The L protein-binding domain (P_L_) is colored magenta. (B) Cryo-EM structure of the L-P complex determined at 3.0 Å resolution. Full-length L protein was incubated with a P protein fragment comprising residues 35–106. Domains are colored as in (A). Flexible P_L_ residues, connecting the three L-bound segments that are not modeled in the structure, are indicated by dashed lines. (C) Close-up view of the cryo-EM density map. The region shows one of two bound zinc atoms. Tetrahedral coordination is with two histidine and two cysteine residues of the Cap.

The polymerase catalytic site faces a cavity at the center of the RdRp domain (Liang et al., 2015). Channels between this cavity and the surface of the molecule allow entry and exit of the template RNA strand and entry of NTPs. A loop from the Cap domain, projecting into the RdRp catalytic cavity, appears to be a priming loop to support the initiating nucleotide. Analogous loops are present in other viral RNA-dependent RNA polymerases that do not initiate on a polynucleotide primer (Lu et al., 2008; Tao et al., 2002). The priming loop must shift out of the way to allow elongation, as seen in the transition from initiation to elongation in the reovirus polymerase (Tao et al., 2002). In the case of VSV L, suitable reconfiguration would allow the leading end of the elongating product strand to move into a cavity in the Cap domain that includes its active-site residues.

The capping reaction catalyzed by the L proteins of non-segmented, negative-sense RNA viruses proceeds through intermediates different from those of the mammalian-cell capping process (and different from the related and well characterized capping mechanism of double-strand RNA viruses). Instead of an attack by the 5’ end of the RNA on a GMP covalently attached to a lysine side chain, the monophosphoylated 5’ end of the transcript attaches covalently to a histidine side chain, and resolution of this intermediate through attack by GDP or GTP results in a guanylated 5’ terminus. The Cap domain is thus a polyribonucleotide transferase (PRNTase). 2’-O methylation and 7-N methylation, both reactions carried out by the single active site on the MT domain, complete the modification.

The resolution of the L-P reconstruction resulted in a few regions of uncertain chain trace and did not allow us to identify features corresponding to P (Liang et al., 2015). It was clear, however, that binding of a P fragment including residues 35-106, the sequence between the conserved N-terminal segment (P_NTD_) and the oligomerization domain (P_OD_) (Figure 1A and Data S2), induced a “closed” structure of L, in which the three terminal domains had well defined positions on the RdRp-Cap structure (Figure 1B). We interpreted the observed conformation as a preinitiation complex. Poor density in the region between the active sites of the RdRp and Cap domains prevented any direct analysis of the coupling of transcription and capping.

We have now extended the resolution of the VSV L-P structure to 3.0 Å, from new images recorded on a Titan Krios microscope. The new reconstruction has allowed us to trace three segments of P bound to L, and to correct a few small errors in the previous model. The molecular L-P interactions explain how P binding leads to a closed structure. The improved definition of several parts of the Cap domain and a clearer trace of the connections into and out of the connector domain have together allowed us to identify a channel leading from the polymerase active site to a potential genome and anti-genome exit site at a position appropriate for acquisition of N, held close to that site by interaction with the N-terminal segment of P. Direct deposition of N onto the emerging RNA could afford prompt encapsidation and maximal protection. Better definition of the Cap active-site cavity also allows us to consider possible conformational changes that accompany formation of the covalent pRNA intermediate and its transfer to GDP.

## RESULTS AND DISCUSSION

### Structure of VSV L-P from Cryo-EM Reconstruction at 3.0 Å Resolution

We prepared the VSV L-P complex for cryo-EM imaging as described previously (Liang et al., 2015), recorded images (Figure S1A and B), and carried out the image processing steps as described in Methods. The final map used for the modeling and interpretation reported here (Figure 1C) had an overall resolution of 3.0 Å as determined by the spatial frequency at which the Fourier shell correlation (FSC) between half maps dropped below 0.143 (Figure S1C). The data quality not only allowed us to improve the overall resolution of the reconstruction, but also to better classify out a more homogenous set of particle images, resulting in a density map in which all domains were equally well resolved.

We could place our previous model (PDB-ID 5A22) in density without any major adjustments. We traced connections into and out of the connector domain more clearly than with the previous, lower resolution reconstruction, corrected a local chain trace error associated with these connections, corrected a few errors in assigning sequence register to segments poorly defined in the previous map, and adjusted many side-chain rotamers, which were generally well defined in the new map. Data collection, model refinement and validation statistics are in Table S1. The positions of various differences between updated and original coordinates are shown in Figure S2 and S3, and listed in Table S2.

### Interactions of the P Subunit with L

The most important new feature was interpretable density for parts of P, enabling us to model three ordered segments from residues 49 to 56, 82 to 89, and 94 to 105 (Figure 2A and B). The density is well defined for those segments (Figure S4). The first segment binds in a shallow pocket on the outward-facing surface of the CTD. The interactions are mostly hydrophobic around a conserved tyrosine residue _P_Tyr53 that also hydrogen-bonds to the side chain of _L_Asp1981 (Figure 2C). P then wraps around the CTD with a stretch of flexible residues that we can only partially see in low-pass filtered density maps (Figure S4B). The second segment binds between the CTD and the RdRp and interacts with the C-terminal arm as it inserts along the edge of the RdRp. Here we observe five hydrogen bonds. On the P side, they all involve main chain atoms, either the amino nitrogen or carbonyl oxygen. These interactions do not directly depend on side chain residues and explain the somewhat relaxed conservation of segment two residues. Binding is further stabilized by hydrophobic contacts of _P_Val84 and _P_Phe87, and a salt-bridge between _P_Glu85 and the CTD residue _L_Lys2022 (Figure 2D). The third segment lies in the groove between the CTD and the connector. We count two salt-bridges and two hydrogen bonds at this binding site. One hydrogen bond, between _P_Glu99 and _L_Gly1199, connects to the CTD. The _P_Tyr95 side chain is stacked between _L_Arg1419 and _L_Tyr1438. A conserved _P_Val102 is accommodated in a hydrophobic L pocked, and a conserved _P_Tyr104 stacks on _L_Pro1535 (Figure 2E). Taken together, the CTD is functionally an adaptor domain of L for its association with P, which then reinforces the interdomain contacts stabilizing the closed conformation we have described.

**Figure 2.**
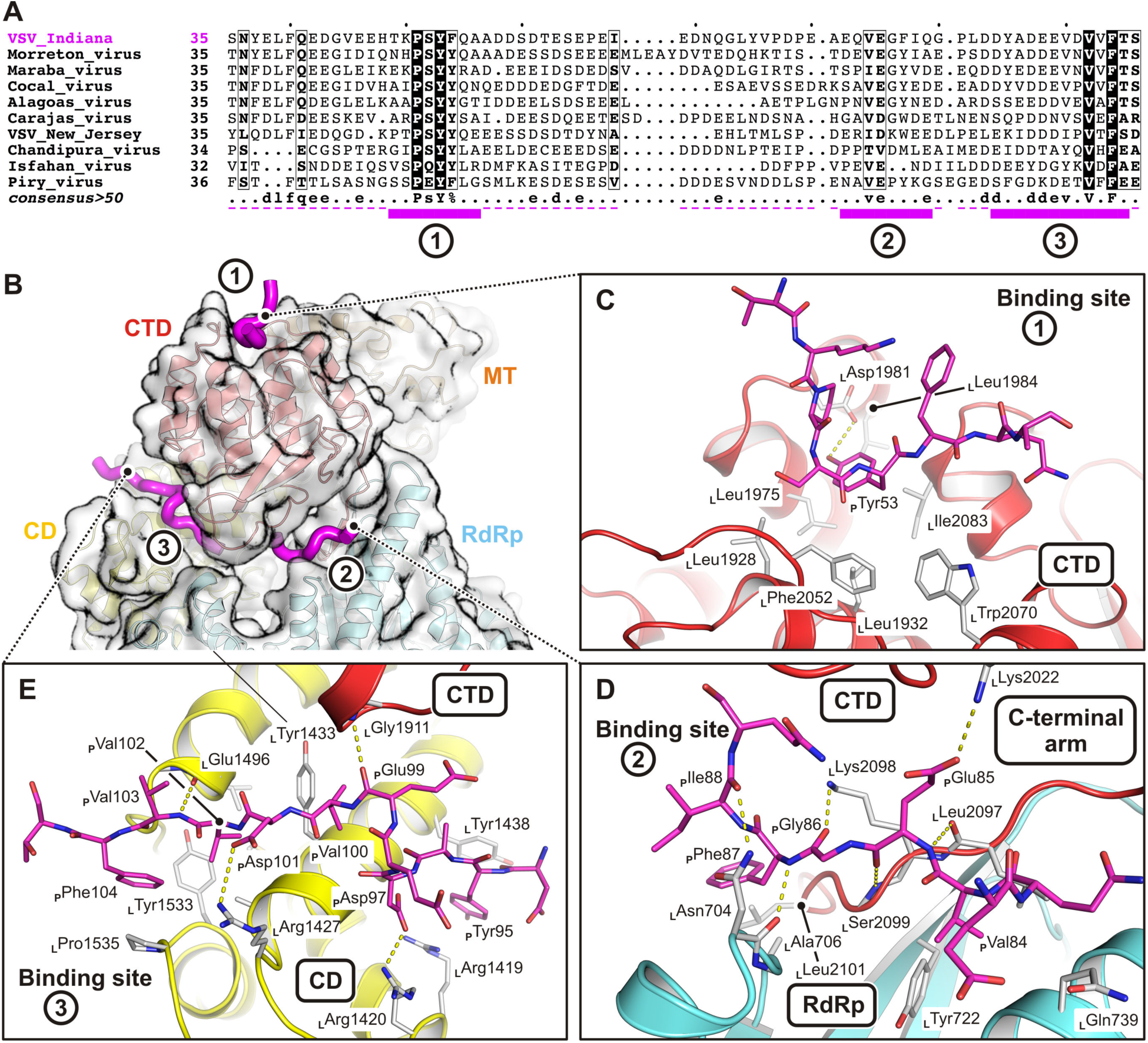
Molecular Details of Phosphoprotein Interactions with the L Protein. (A) Multiple sequence alignment of phosphoprotein amino acid sequences from different viruses. For sequence accession identifiers see Data S2. The alignment is shown for VSV phosphoprotein residues included in the expression construct only. Magenta solid bars at the bottom of the sequences are residues modeled in our structure, magenta dashed lines are residues included in the expression construct. Binding motifs 1, 2, and 3 are labeled. (B) Overview of phosophprotein binding to the L protein. The P protein is shown as a magenta tube. The L protein is colored as in Figure 1. Motif 1 binds on the outer surface of the CTD. Motif 2 binds between the CTD and the RdRp, and also interacts with the L protein C-terminal arm. Motif 3 interacts with the CD and the CTD. (C, D and E) Molecular interactions of phosphorprotein motifs 1 (A), 2 (B) and 3 (C) with the L protein, respectively. The L protein backbone is shown in ribbon representation and colored as in Figure 1. Interacting residues (main chain and/or side chain atoms) are shown as gray sticks. P protein residues are shown as magenta sticks. Hydrogen bonds and salt bridges are indicated with yellow dashed lines.

VSV P dimerizes through a domain (P_OD_) that extends from residue 109 to residue 170 (Figure 1 and Data S2). We analyzed, by projection matching with the present structure (Figure S5 and Data S3), images of the negatively stained L-P dimers described in much earlier work (Figure 3A) (Rahmeh et al., 2010). We found that the mean distance between residues 105 on the P chains of the two matched complexes (60 Å) (Figure 3B) agreed very well with what we expect from the known structure of the dimerization domain (Ding et al., 2006), allowing for extension of the short segment from 105 to 109 on each of the dimerized complexes (Figure 3C). The good agreement validates the polarity of the P chain trace and the assignment of residues in segment 3.

**Figure 3.**
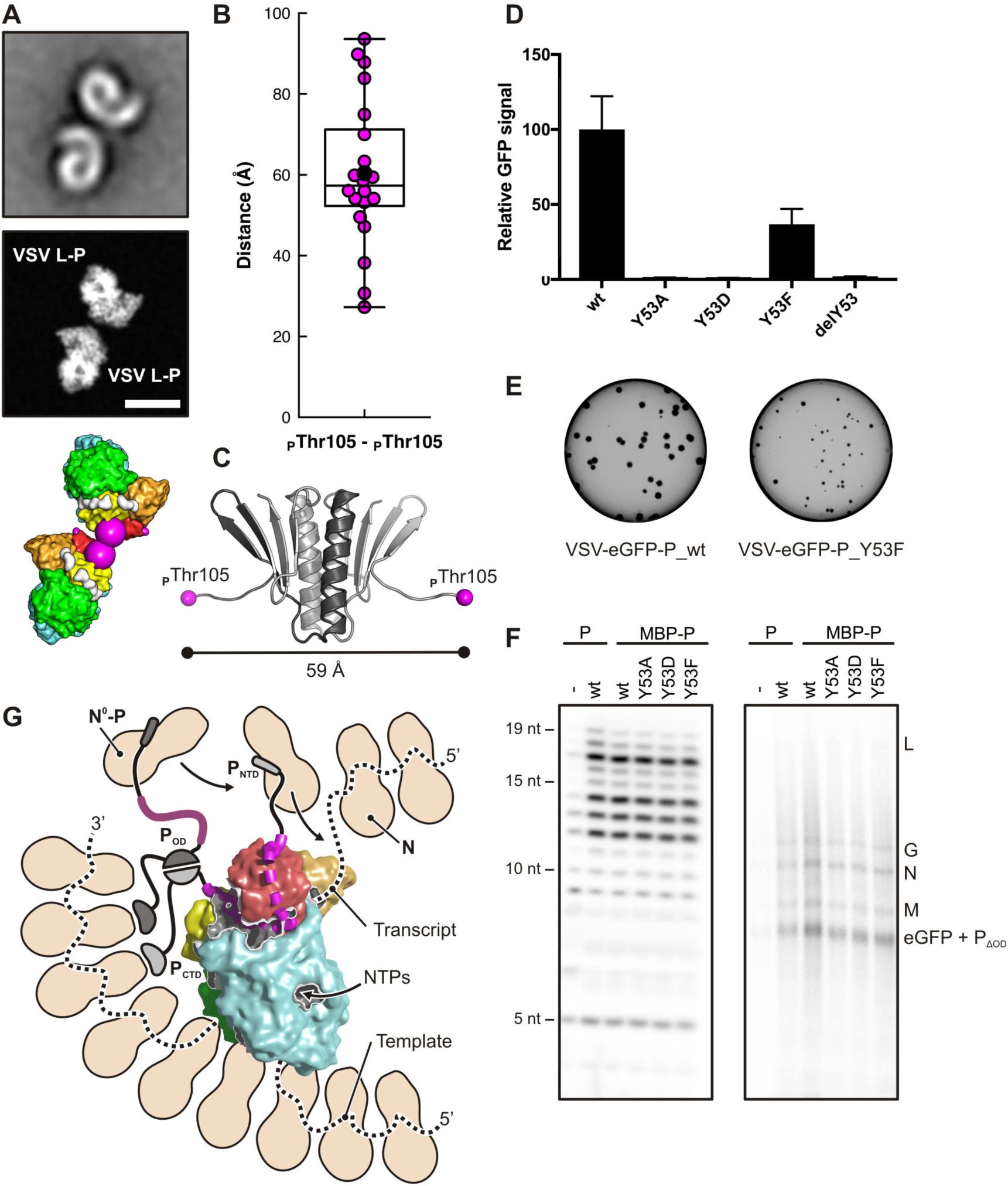
Characterization of Phosphoprotein Binding and Function. (A) Projection matching of VSV L-P dimer negative-stain class averages. Top, representative negative-stain class average (Rahmeh et al., 2010). Middle, projection image of our reconstruction obtained from two aligned protomers. The scale bar corresponds to 100 Å. Bottom, two aligned protomers of the L-P structure shown in surface representation. The last modeled P protein residues (_P_Thr105) are shown as large magenta spheres. See Figure S5 and Data S3 for details of the projection matching analysis. (B) Statistical analysis of the distance between P protein binding motif 3 C-terminal residues (_P_Thr105) in the negative-stain class averages of VSV L-P dimers. For each of the 20 class averages the distance obtained from the projection matching analysis is shown as a magenta dot. The average distance is shown as a black dot. (C) Structural model of the P protein oligomerization domain (P_OD_). The dimer structure is from PDB-ID 2FQM (residues 111–171) (Ding et al., 2006); residues 105–110 were modeled in extended conformation. The distance between _P_Thr105 residues is 59 Å. (D) Cell-based assay using a eGFP reporter as a readout for L activity when RNA synthesis is reconstituted with the various P mutants. (E) Plaque assay with VSV-eGFP-P_wt and VSV-eGFP-P_Y53F. GFP signal was monitored 24 hours post-infection. (F) Reconstituted RNA synthesis in vitro on naked RNA (left) and an N-RNA template (right). (G) Model for replication of a ribonucleoprotein (RNP) complex by VSV L-P. The N protein is colored beige. The RNA template accesses the RdRp active site after dissociation from about three N subunit of the RNP. P binding stabilized the closed conformation of L, leads to processive polymerization, and positions the N-terminal domain (P_NTD_) close to a putative product exit channel to encapsidate the emerging RNA by transferring N from N^0^-P. The C-terminal domain of P (P_CTD_) can interact with the RNP and may guide the polymerase along the template. Dimerization of P through the oligomerization domain (P_OD_) would lead to additional RNP and N^0^-P interactions.

In segment 1, tryosine _P_Tyr53 fits snugly into a pocket on the outward-facing surface of the CTD. For a series of mutations at position 53, we found that the mutants greatly decreased RNA synthesis in cells transfected with L, P, N and infected with particles expressing a reporter template (Figure 3D). When incorporated into virions, the mutations resulted in no growth (Y53A and Y53D) or substantially attenuated growth (Y53F) (Figure 3E). They did not, however, affect *in vitro* transcription by L-P (Figure 3F). Anchoring of the P N-terminus at the apex of the CTD (Figure 3G) thus contributes to RNA synthesis during viral infection but does not influence the RNA polymerase activity directly. The most plausible interpretation of these results is that the defect is in replication, perhaps because of failure to deliver N to the nascent RNA.

### Priming, Initiation, and Elongation

The loop between residues 1152 and 1173 in the Cap domain projects into the catalytic cavity of the RdRp and appears to be a priming loop to support the nucleotide that initiates polymerization. As shown in Figure 4A, superposition of the catalytic residues of L on corresponding residues of the reovirus λ3 initiation complex (Tao et al., 2002) (Table S3 and Figure S6) places the initiating nucleotide so that it stacks directly on a prominent tryptophan (_L_Trp1167) at the tip of the VSV L priming loop. Mutation of this tryptophan to alanine severely restricts initiation at the end of the genome or antigenome; mutation to tyrosine or phenylalanine has a smaller effect (Ogino et al., 2019). Mutation of the homologous tryptophan in rabies virus (RABV) L also compromises end initiation, but it appears not to affect internal initiation of transcripts, nor does it prevent capping (Ogino et al., 2019).

**Figure 4.**
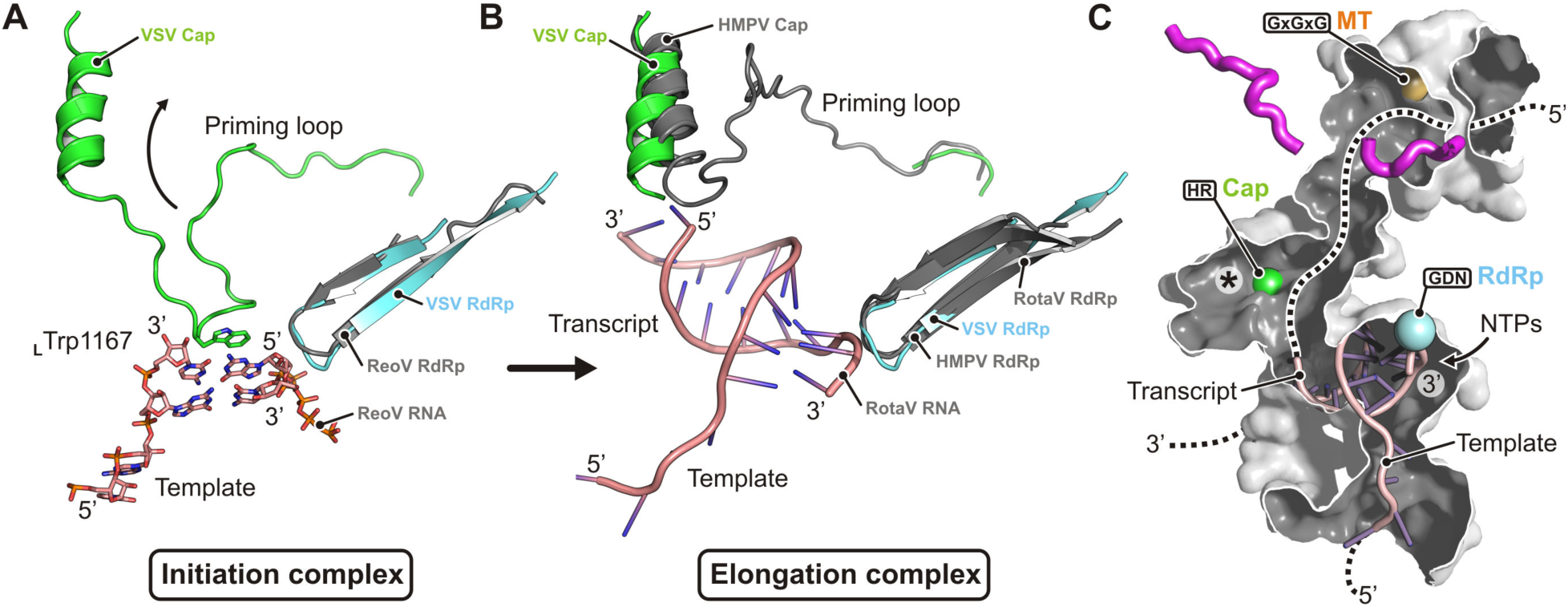
Model for RNA Synthesis by the VSV L-P Complex. (A) Model of an initiation complex. The VSV L protein priming loop of the Cap is shown in green, the RdRp hairpin harboring the catalytic GDN motif is shown in cyan. Template and transcript nucleotides are from the crystal structure of the reovirus λ3 polymerase initiation complex (ReoV) (PDB-ID 1N1H) (Tao et al., 2002) superimposed on the VSV L-P structure. The first phosphodiester bond of the transcript is formed between the nucleotide in the priming position and the incoming nucleotide. _L_Trp1167 supports the priming nucleotide by stacking interaction with the base. (B) Model of an elongation complex. To allow passage of the growing template-transcript RNA duplex, shown here from the superimposed structure of transcribing rotavirus VP1 polymerase (RotaV) (PDB-ID 6OJ6) (Jenni et al., 2019), the priming loop may retract to a position similar to the one observed in the structure of the human metapneumovirus polymerase (HMPV) (PDB-ID 6U5O). (C) Tunnels in the VSV L-P structure. The tunnel is cut open to allow view of the central cavities. The RNA template and transcript strands are from the rotavirus structure as in (B). The template-entry channel is at the bottom. The template-exit channel is at the back, bottom-left. Nucleotide substrates enter the catalytic site laterally from the right side (shown by an arrow). Catalytic sites with conserved residues for polymerization (RdRp), capping (Cap) and methylation (MT) are show by colored spheres. The tunnel was probed after omitting priming loop residues 1152–1173. Priming loop retraction into the space between the Cap and CD domains (shown by an asterisk) opens a continuous path for the transcript RNA (shown by a dashed line).

Further polymerization requires retraction of the priming loop or, potentially, displacement of the entire Cap domain. Recently determined structures of the human metapneumovirus (HMPV) (Pan et al., submitted) and the respiratory syncytial virus (RSV) (Gilman et al., 2019) polymerases show a fully retracted priming loop, with no substantial displacement of the rest of the Cap domain. The priming loops of the two enzymes and the catalytic cavities of the Cap domains are very similar, and we can show by superposition that the VSV priming loop can also retract without any major rearrangements in the rest of the Cap domain (Figure 4B). Moreover, when we model the VSV L priming loop based on the HMPV L protein structure, we find that a twisting tunnel connects the RdRp catalytic site with an opening where the CTD, RdRp and MT all meet (Figure 4C). We suggest that this tunnel might be the exit pathway for uncapped products – i.e., the full-length antigenome and genome during replication and the uncapped leader during transcription –, which bypass the active sites of the Cap and MT domains.

The N-terminal segment of P (P_NTD_) binds a single subunit of N, forming the so-called N^0^-P complex. The likely function of this complex is to deliver N to the nascent genome and antigenome as they emerge from the polymerase. About 15 residues connect the P-binding element (residues 6-34) with the segment of P docked onto the outer face of the CTD. The N^0^ passenger could therefore lie close to the proposed exit site for upcapped products. A possible explanation for the defect in Y53 mutants is release of the docked N-terminal region of P from its site on the CTD, causing inefficient delivery of N to the replication products. This defect would not affect transcription.

At the other end of the P polypeptide chain, a small C-terminal domain (P_CTD_) interacts with RNP. The binding site is at the C-terminal lobe of the N proteins (Green and Luo, 2009). Together with a presumptive interaction of RNP with L at the template entry site, the contact with P could facilitate passage of the template strand through the enzyme. We have proposed previously that the subunits of N (probably three) that need to separate from the RNA template as it threads through the polymerase do not dissociate from each other and instead processively rebind the RNA as polymerization proceeds. The N-N interaction creates a chain-like array, with an N-terminal arm and a C-proximal loop contacting the two neighboring subunits in the chain (Ge et al., 2010; Green et al., 2006). Interaction with the C-terminus of L-bound P could then ensure correct reassociation of this putative chain of N subunits with exiting RNA (see Figure 3G).

Transcription of all products except the uncapped leader requires a different sequence of events. One possibility is that the immediately upstream transcript leaves and the priming loop snaps back into the position seen in our current structure. Internal initiation does not appear to depend on _L_Trp1167, however, and another possibility is that the 3’ end of the departing transcript, still base paired with a few residues of the oligo-U tract at the 3’ end of the upstream gene, supports the initiating NTP instead. The interaction would need to take the intergenic GA sequence on the template into account. Further polymerization would then displace the upstream transcript (Shuman, 1997).

Acquisition of the guanylate cap occurs when the transcript length has reached at least 31 nucleotides (Tekes et al., 2011). If we suppose that formation of the covalent intermediate with histidine _L_His1227 occurs before addition of the thirty-first nucleotide to the transcript, then approximately 30 nucleotides of transcript would need to fit into the combined active-site cavities of RdRp and Cap. The volume of the Cap active-site cavity, marked with an asterisk in Figure 4C, is 1.2×10^4^ Å^3^. Assuming a volume occupied per Dalton (Da) of RNA (V_m_) of 2.1 to 4.6 Å^3^ and an average mass of 340 Da per nucleotide (Speir and Johnson, 2012), the interior of the Cap domain could accommodate between 8 and 17 transcript nucleotides, in addition to the ∼10 nucleotides still based-paired with template in the RdRp catalytic cavity. These relatively generous estimates suggest that addition of the guanylate to the 5’ end of the transcript requires opening of the multi-domain assembly from the closed conformation present in our structure. A plausible transition would be release of the CD, MT and CTD from their docked positions, as indeed occurs in the absence of P. If that transition also enabled a guanine nucleotide to access the RNA-protein covalent linkage, release of the capped 5’ end would allow it to diffuse efficiently into the MT active site, since both the transcript (still anchored to the RdRp at its growing, 3’ end) and the MT (linked to RdRp-Cap through the CD) cannot move far from each other.

## STAR METHODS

### Cryo-EM Structure Determination

The VSV L-P protein complex was prepared for structural analysis as previously described (Liang et al., 2015). Briefly, 3 µL of protein at ∼0.35 mg/mL were applied to a C-flat grid (CF-1.2/1.3-4C, Protochips) that had been glow discharged at 40 mA for 30 s. Grids were plunge-frozen with an Vitrobot Mark II (FEI Company) after blotting for 3 s at 4°C and ∼85% relative humidity. 3307 movies of vitrified protein solution were collected using SerialEM (Mastronarde, 2005) on a Titan Krios electron microscope (ThermoFisher) at 300 kV and 59,000 nominal magnification, on a K2 Summit detector (Gatan, Inc.) operated in counting mode, resulting in a calibrated pixel size of 0.485 Å. Each movie contained 50 frames, collected in a 10 s exposure at 2 electrons/Å^2^/frame. Most micrographs showed well-dispersed particles (Figure S1A). Movies were aligned and exposure-filtered using Unblur (Grant and Grigorieff, 2015). 3235 movies were selected for further processing based on the frame motion traces, excluding movies with traces suggesting large and zig-zag motion. The contrast transfer function (CTF) was determined using CTFFIND4 (Rohou and Grigorieff, 2015). CTF fringes on many micrographs were fitted to better than 3.0 Å resolution (Figure S1B). 188,004 particles were picked with IMAGIC (van Heel et al., 1996), command PICK_M_ALL. Particles were windowed into 400×400 pixel images and aligned using *cis*TEM (Grant et al., 2018) against the pixel size-adjusted 2015 map (EMD-6337) at an initial resolution of 8.0 Å, then refined at successively higher resolution, up to a resolution limit of 5.5 Å. Refinement was continued with 3D classification of two classes, at increasing resolution up to 4.0 Å, resulting in two reconstructions at 3.6 and 3.2 Å resolution (69,312 and 118,692 particles, respectively). Particles assigned to the 3.2-Å reconstruction were further refined at 4.0-Å resolution, and classified again into two classes with reconstructions at 3.8 and 3.1 Å resolution (34,713 and 83,979 particles, respectively). Further refinement of the best class at 4.0 Å resolution resulted in the final 3.0 Å reconstruction. The reconstruction was sharpened using *cis*TEM’s sharpening tool, amplifying signal at high resolution with a *B* factor-equivalent of 75 Å^2^.

### Model Building and Refinement

We used the program O (Jones et al., 1991) to adjust our previously determined VSV L protein structure (PDB-ID 5A22, determined at 3.8 Å resolution) (Liang et al., 2015) to fit the 3.0 Å resolution cryo-EM map. We modeled the P protein residues *de novo*. We refined coordinates and *B* factors with phenix.real_space_refine (version 1.16-3549) (Afonine et al., 2018). In addition to the standard chemical and grouped *B* factor restraints, we used secondary structure, rotamer, and Ramachandran restraints (weight=1.0, nonbonded_weight=500, rama_weight=6.0). We used phenix.mtriage (Afonine, 2017) and MolProbity (Chen et al., 2010) to calculate the validation results shown in Figure S1C and Table S1. The final VSV L-P model comprises L protein residues 35–1211, 1217–1332, 1339–1590, and 1596–2109; P protein residues 49–56, 82–89, and 94–105; and two zinc atoms.

### Projection Matching of Negative-Stain Class Averages

We extracted 20 negative-stain class averages of L-P dimers from a screenshot of a previously published figure (Fig. S4C) in (Rahmeh et al., 2010) using RELION-3 (Zivanov et al., 2018). From those dimer images we manually extracted 40 subparticles, one of each protomer, and masked them with a soft spherical mask. We obtained rotation angles for reference projections of our reconstruction with healpy (http://healpy.readthedocs.org), the Python implementation of the HEALPix library of routines for equal area sampling of a sphere (Górski et al., 2005). Because the L-P complex has no symmetry and we did not allow mirroring of reference projections in the subsequent alignment step, we sampled angles on the entire sphere. We aligned the subparticle images to the reference projections with e2simmx.py from EMAN2 (Bell et al., 2016). Alignments were scored by calculating a cross-correlation coefficient (CCC). Because information in the negative-stain class averages is restricted to low resolution, reference projections from very different angles resulted in similar CCCs, and the “correct” solution was not always the top scoring projection. We therefore looked at the top five alignment peaks, which we located with the CCP4 program PEAKMAX (Winn et al., 2011), and then manually selected for further analysis the one whose reference projections most closely matched the particle image as judged by eye (Figure S5). To calculate the distance between the P protein C termini of the two promoters, we transformed the model coordinates according to the determined alignment parameters. A representative example of the analysis results for class average 1 is shown in Figure S5. The analysis results for all class averages can be found in Data S3.

### Functional Assays

#### Cells

BSR-T7 cells (a kind gift from K. Conzelmann) (Buchholz et al., 1999) were maintained in Dulbecco’s modified Eagle’s medium (DMEM; Corning Inc., 10-013-CV) containing 10% fetal bovine serum (FBS; Tissue Culture Biologicals, TCB 101) at 37 °C and 5% CO_2_.

#### Plasmids

For mammalian expression, plasmids expressing Y53 mutants were generated by site-directed mutagenesis using the Q5 polymerase (New England Biolabs) on the parental P expressing plasmid (pMB-NS) (Pattnaik and Wertz, 1990) with dedicated primers (for Y53A: TAGGCCCTCTGCGTTTCAGGCAGC and GTATGCTCTTCCACTCCG, for Y53D: TAGGCCCTCTGATTTTCAGGC and GTATGCTCTTCCACTCCG, for Y53F: CATACTAGGCCCTCTTTTTTTCAGGCAGC and AGAGGGCCTAGTATGCTC, for deltaY53: TTTCAGGCAGCAGATGATTC and AGAGGGCCTAGTATGCTC). For bacterial expression and purification, fusion proteins made of a 6 histidine tag followed by the maltose-binding protein (MBP), the human rhinovirus type 14 (HRV) 3C protease cleavage site and the P protein, were cloned into a pET16b plasmid. The MBP tag was added to increase P solubility. A PCR product containing the 6xHis-MBP fragment was amplified from pET16b-MBP-MACV-Z (Kranzusch and Whelan, 2011) using AAGGAGATATACATATGGCTCACCATCACCATCACC and CTGGAACAGTACTTCCAAACCAGAACCCTCGATCCCGAGGTTG, and one containing the HRV_3C-P fragment was amplified from pET16b-P (Rahmeh et al., 2012) using GAAGTACTGTTCCAGGGTCCTATGGATAATCTCACAAAAGTTC and GTTAGCAGCCGGATCCA. Both were inserted into pET16b-MBP-MACV-Z digested with NdeI and BamHI using the Infusion kit (Takara) to form pET16-6xHis-MBP-3Cc-P. Y53 mutations were introduced in pET16-6xHis-MBP-3Cc-P by site-directed mutagenesis with primers described above. To rescue viruses expressing P_Y53 mutants, Y53 mutations were introduced into pVSV1(+)-eGFP backbone (Whelan et al., 2000).

#### Protein purification

L and P proteins were purified as previously described (Rahmeh et al., 2012). MBP-P were purified from Rosetta cells (EMD Millipore) after induction with 0.3 mM IPTG overnight at 18 °C. Cells were then pelleted and resuspended with 5 mL cold lysis buffer per gram of pellet (50 mM HEPES (pH 7.4), 100 mM NaCl, 5% glycerol, 5 mM imidazole, 5 mM 2-mercaptoethanol and 1X protease inhibitor cocktail [cOmpleteTM, Roche Cat. # 4693116001]). After sonication, 40 mL cell lysate were clarified by centrifugation at 20,000 x g for 20 min, and the supernatant was incubated with 2 mL of Ni-NTA beads (Qiagen) overnight at 4 °C. Beads were washed three times with lysis buffer (without protease inhibitor), and MBP-P proteins were eluted in 5 mL elution buffer (50 mM HEPES (pH 7.4), 100 mM NaCl, 5% glycerol, 5 mM imidazole and 5 mM 2-mercaptoethanol). The eluate was concentrated with a centrifugal filter (Amicon) and passed through a Superdex S200 size exclusion chromatography column (GE Healthcare) in storage buffer (20 mM HEPES (pH7.4), 100 mM NaCl, 5% glycerol and 5 mM 2-mercaptoethanol). MBP-P proteins eluted in a single peak and were kept at −80 °C in storage buffer.

#### Virus

Attempts to rescue viruses expressing P_Y53 mutants, were performed as previously described (Whelan et al., 1995) using pVSV1(+)-eGFP-P_Y53x plasmids. VSV-eGFP-P_Y53F was the only mutant virus forming plaques. It was amplified and titered on BSR-T7 cells.

#### N-RNA purification

Nucleocapsid template was purified essentially as described previously (Ongradi et al., 1985). To limit background activity due to residual P bound to N-RNA, nucleocapsids were extracted from a recombinant VSV expressing a monomeric P protein whose oligomerization domain has been deleted (VSV-P_ΔOD_). Briefly, 10 mg of gradient purified virions were disrupted on ice for 1 h in 10 mL of virion disruption buffer (VDB: 20 mM Tris-HCl (pH 7.4), 0.1% Triton X-100, 5% glycerol, 1 mM EDTA, 2 mM dithioerythritol and 600 mM NH_4_Cl). Lysed virions were centrifuged at 240,000 × g for 3.5 h through a glycerol step gradient of 0.25 ml each of 40, 45, and 50% glycerol in VDB. Pellets were resuspended overnight in 0.5 ml of 10 mM Tris-HCl (pH 7.4), 100 mM NaCl, 1 mM EDTA and 1 mM DTT, and disrupted again on ice for 1 h after mixing with 0.5 mL of 2X high salt VDB (1X HSVDB: 20 mM Tris-HCl (pH 7.4), 0.1% Triton X-100, 5% glycerol, 1 mM EDTA, 2 mM dithioerythritol and 1.5 M NH_4_Cl). Nucleocapsids were isolated by banding on a CsCl gradient. In a 5 mL tube, 1 mL of disrupted virions was added at the top of a step gradient of 2 mL of each 20 and 40% (wt/wt) CsCl in HSVDB. After centrifugation at 190,000 × g for 3 h, N-RNA were recovered by side puncture and diluted fivefold in NaCl-Tris-EDTA-DTT buffer (NTED: 10 mM Tris-HCl (pH 7.4), 100 mM NaCl, 1 mM EDTA and 2 mM DTT). N-RNA was then centrifuged through a 0.5-ml cushion of 50% glycerol in NTED buffer, resuspended overnight in NTE buffer and stored in aliquots at −80°C.

#### Production of minigenome particles

Production of transcription competent minigenome particles was as described previously with minor modifications (Pattnaik and Wertz, 1990; Wertz et al., 1994). BSR-T7 cells were plated in a 60mm dish and infected the next day with a vaccinia virus expressing T7 polymerase (vTF7-3) (Fuerst et al., 1986) at MOI 3 for 1 h in Dulbeco’s Phosphate Buffered Saline liquid (DPBS; Sigma Cat# 59300C). Cells were then transfected using Lipofectamine 2000 with plasmids expressing N (5.5 µg), P (1.6 µg), M (1.25 µg), G (2.5 µg), L (0.5 µg) and a minigenome containing an eGFP reporter gene (7.5 µg). Four hours later, the medium was replaced with 3 mL DMEM containing 2% FBS. The cell supernatant containing the DI particles (DI stock) was harvested 2 days after transfection.

#### Cell-based gene expression assay

BSR-T7 cells were plated in a 96-well plate and infected the next day with a vaccinia virus expressing T7 polymerase (vTF7-3) at MOI (multiplicity of infection) 3 for 1 h in DPBS. Cells were then transfected with plasmids expressing N (91 ng), P (26 ng) and L (7.5 ng) using Lipofectamine 2000. Four hours later, cells in each well were infected with 3 µL DI stock in 30 µL DMEM for 1 h. Two days after transfection, GFP signal was measured using a Typhoon FLA 9500 scanner (GE Healthcare).

#### *In vitro* transcription assays

RNA synthesis assay on naked RNA was described previously (Morin et al., 2012). Briefly, purified L (100 nM) was incubated with purified P or MBP-P (250 nM) and a 19nt-long RNA corresponding to the first nucleotides of the genomic promoter (1.4 µM). Reactions were performed in 10 µL at 30 °C for 5 h in presence of radiolabeled GTP (^32^P-αGTP, PerkinElmer) and analyzed on a 20% polyacrylamide/7M urea gel. The gel was exposed overnight to a phosphor screen (GE Healthcare), which was scanned on a Typhoon FLA 9500 scanner. The transcription assay on encapsidated template was carried out as described elsewhere (Li et al., 2008) by incubating 1 µg of purified L protein with or without 0.5 µg of P or 1.2 µg of MBP-P proteins and 5 µg of N-RNA purified from VSV-P_ΔOD_ virions. Reactions were performed in 100 µL in the presence of 1 mM ATP, 0.5 mM CTP, GTP, and UTP, 15 µCi of radiolabeled GTP (^32^P-αGTP, PerkinElmer), 30% vol/vol rabbit reticulocyte lysate (Promega), 0.05 µg/µL actinomycin D (Sigma) in transcription buffer (30 mM Tris-HCl pH 8.0, 33 mM NH_4_HCl, 50 mM KCl, 4.5 mM Mg(OAC)_2_, 1 mM DTT, 0.2 mM spermidine and 0.05% Triton X-100) at 30 °C for 5 h. RNA was extracted with Trizol reagent (Invitrogen Cat# 15596018) following manufacturer’s protocol, boiled at 100 °C for 1 min, incubated on ice for 2 min, mixed with a 1.33 X loading buffer (33.3 mM citrate pH 3, 8 M urea, 20% sucrose and 0.001% bromophenol blue) and analyzed on a 25 mM citrate pH 3, 1.75% agarose, 6 M urea gel run for 18h at 4 °C and 180 V. Gels were fixed (in 30% methanol and 10% acetic acid), dried, and exposed overnight to a phosphor screen (GE Healthcare), which was scanned on a Typhoon FLA 9500 scanner.

### Superposition of Atomic Models

To compare the VSV L-P structure to other viral polymerases, we used the program LSQMAN (Kleywegt and Jones, 1997) for superposition, except for the structure of the reovirus λ3 polymerase initiation complex, for which we also used the program O (Jones et al., 1991) to focus the alignment on the RdRp active site. Superposition statistics are summarized in Table S3, and superposed models are shown in Figure S6.

### Cavity Volume Calculation

We used the program VOIDOO (Kleywegt and Jones, 1994) to calculate the volume of the Cap active site cavity (Figure 4C, marked by and asterisk). The radius of the probe sphere was 1.4 Å. The calculated volume corresponds to the probe-occupied volume. We closed off the cavity of interest at the constrictions leading to the RdRp and MT active sites cavities, respectively, with dummy atoms.

### Figure Preparation

We used MAFFT to calculate the amino acid multiple sequence alignments (Katoh et al., 2002). We prepared the figures with PyMol (The PyMOL Molecular Graphics System, Version 2.1 Schrödinger, LLC), Pov-Ray (www.povray.org), matplotlib (Hunter, 2007), and ESPript (Robert and Gouet, 2014).

### Accession Identifiers of Density Maps and Atomic Coordinates

We have deposited the cryo-EM maps in the Electron Microscopy Data Bank with accession identifier EMD-20614 and the atomic coordinates of the VSV L-P complex in the Protein Data Bank with accession identifier PDB-ID 6U1X.

## SUPPLEMENTAL INFORMATION

Supplemental Information can be found online.

## ACKNOWLEDGMENTS

We acknowledge support from NIH grant R37 AI059371 to S.P.J.W. N.G. and S.C.H. are Investigators in the Howard Hughes Medical Institute.

---

**Figure S1.**
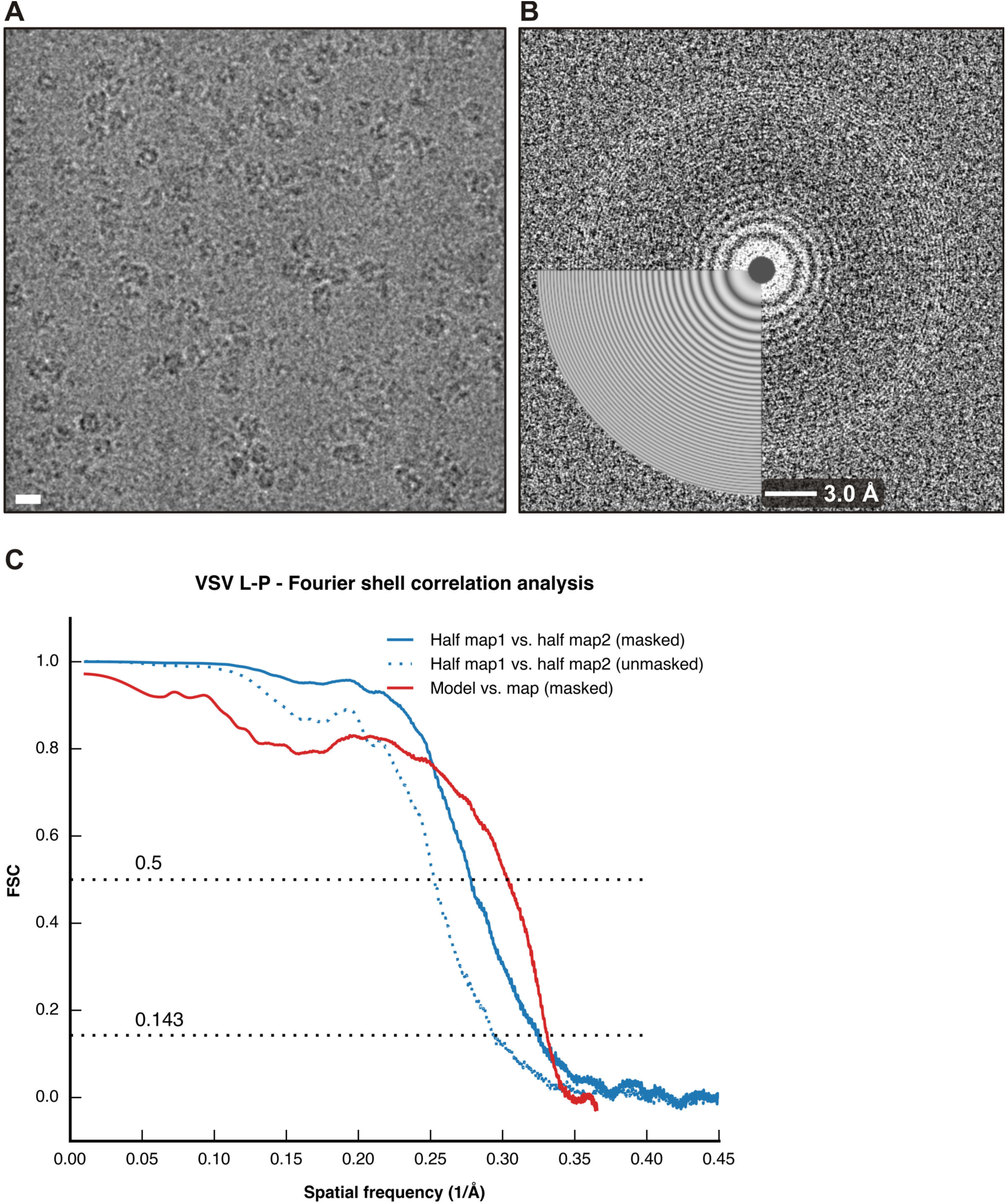
Fourier Shell Correlation (FSC). Related to STAR Methods. (A) Micrograph of vitrified VSV L-P, showing well-dispersed particles. The scale bar corresponds to 100 Å. (B) Contrast transfer function (CTF) fringes calculated from the movie frames of the micrograph shown in A, with good match of the fitted pattern. (C) FSC curves for the VSV L-P cryo-EM reconstruction and refined model calculated with phenix.mtriage (Afonine, 2017). Correlations for the two half maps are shown in blue after applying a loose spherical mask (dashed line) and a tight mask based on the model (solid line), respectively. The correlation between the refined model and final map is shown as red solid line.

**Figure S2.**
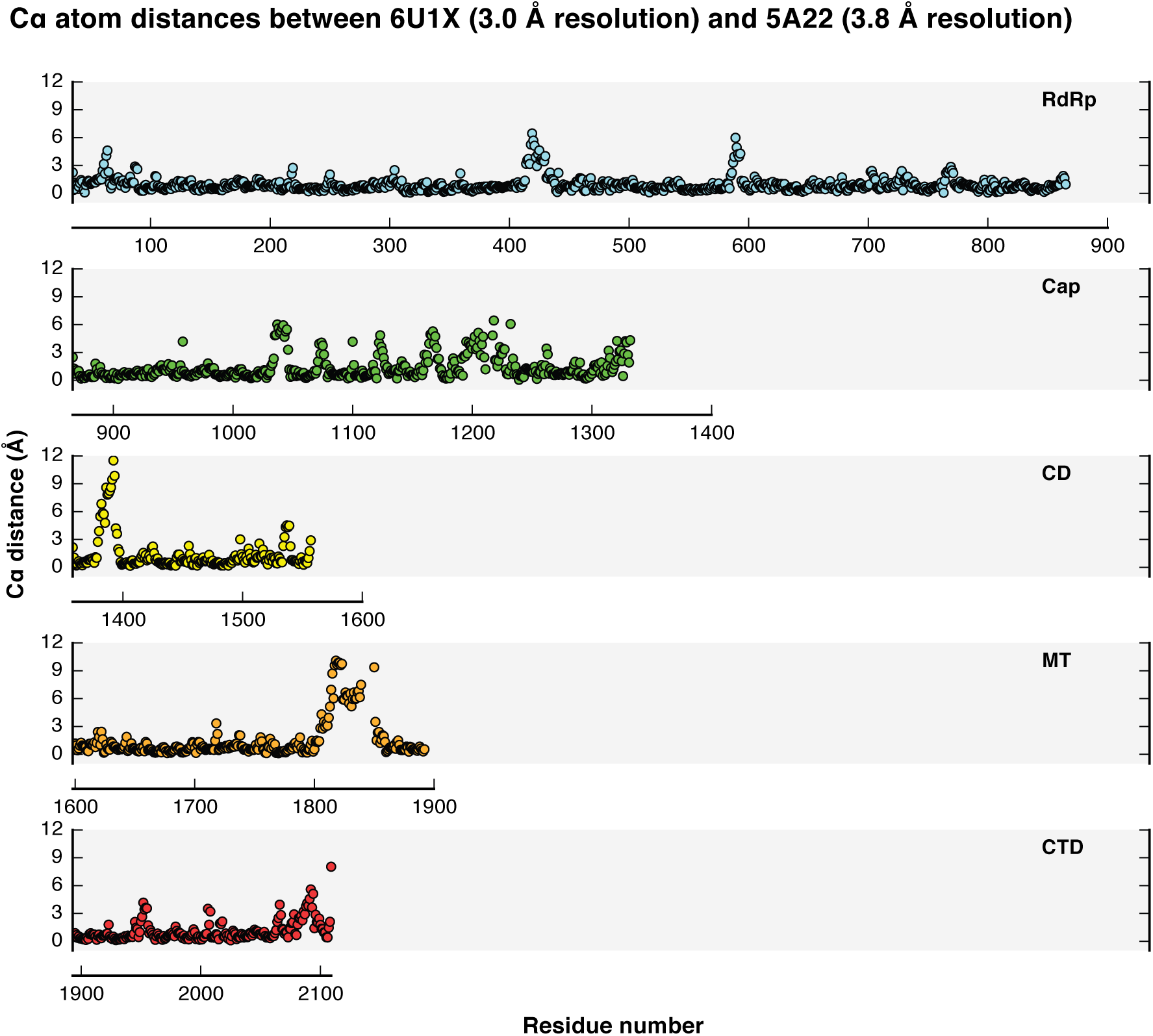
Cα Distances between PDB-ID 6U1X and PDB-ID 5A22. Related to Figure 1 and STAR Methods. The plot shows the Cα distances for each of the L protein domains after superposing the 3.0 Å resolution structure reported here (PDB-ID 6U1X) onto our previously determined structure at 3.8 Å resolution (PDB-ID 5A22). Domains are colored as in Figure 1. See also Table S2 for regions with most substantial modeling improvements.

**Figure S3.**
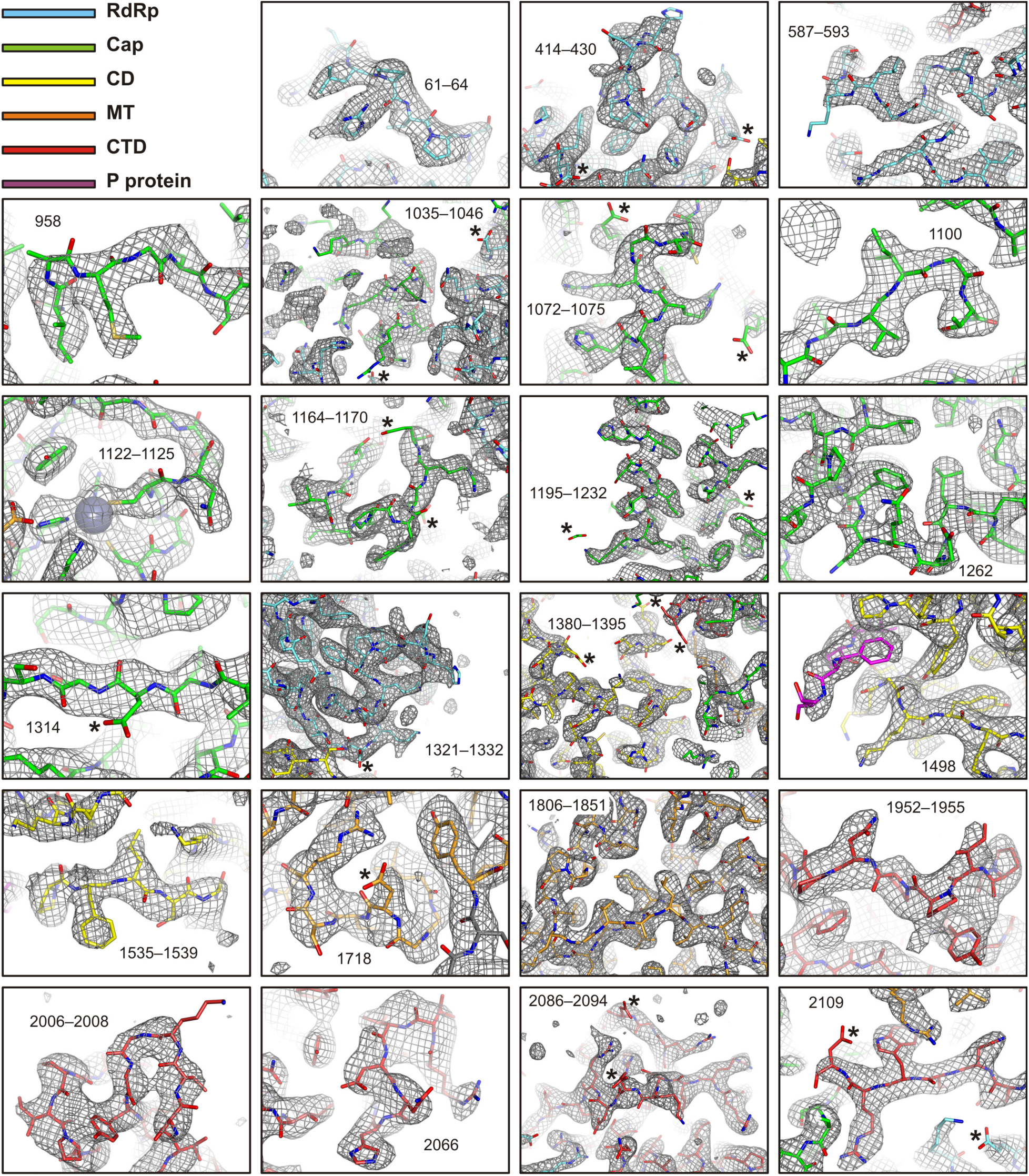
Cryo-EM Density of Improved Model Regions of the VSV L-P Complex. Related to Figure 1 and STAR Methods. The density map (EMD-20614, B sharpened map) is shown as gray mesh. Carbon atoms of the protein domains are colored as in Figure 1. Nitrogen atoms are blue; oxygen, red; sulfur, orange. Acidic amino acid side chains (Asp and Glu) with characteristic absence of density in the cryo-EM reconstruction are labeled with asterisks.

**Figure S4.**
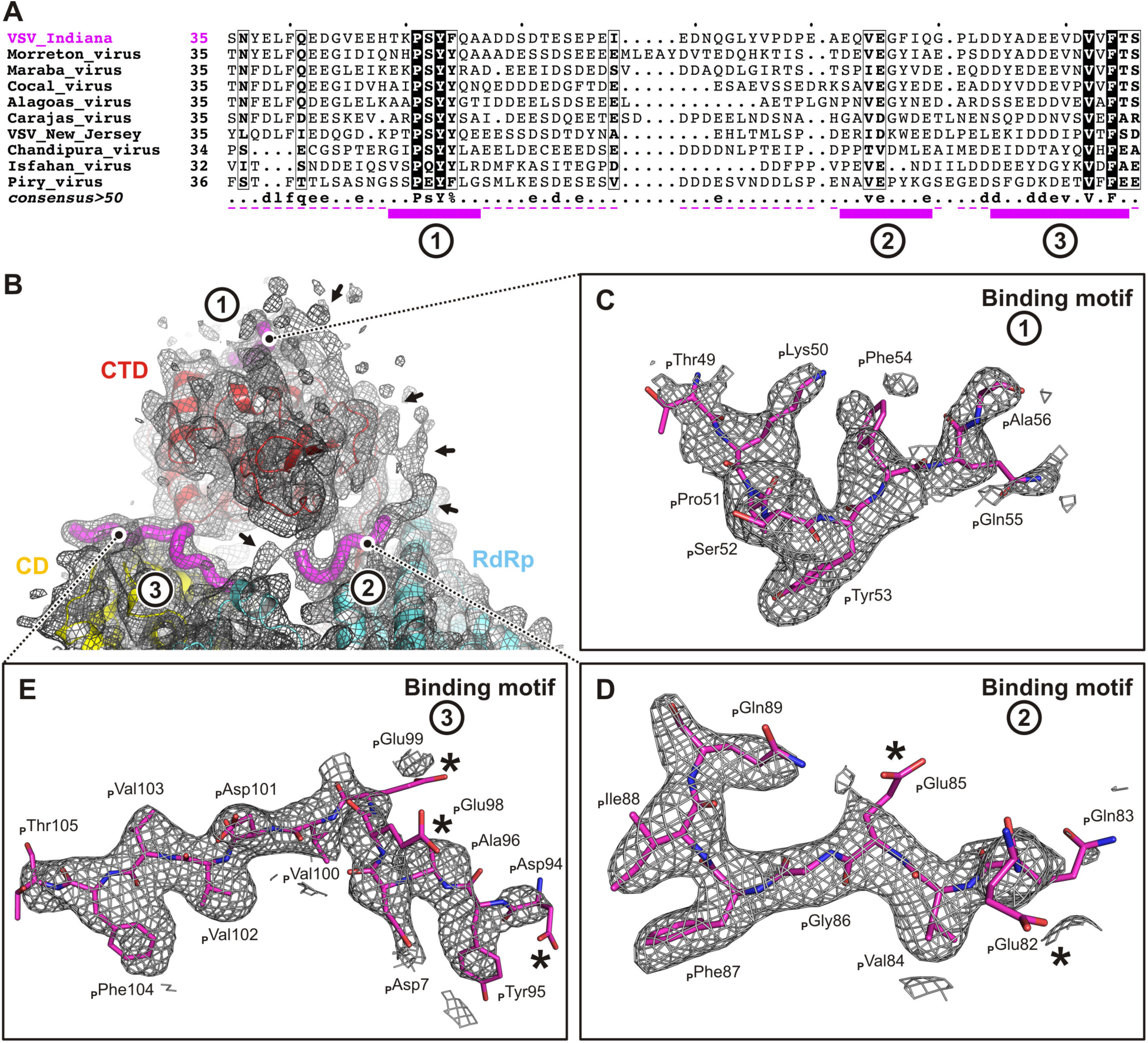
Cryo-EM Density of the Phosphoprotein. Related to Figure 2. (A) Multiple sequence alignment as shown and described in Figure 2A. (B) A low-pass filtered map to 5 Å resolution of the VSV L-P reconstruction shows additional density for some less well ordered segments of the P protein (highlighted with arrows), corresponding to residues connecting the L protein-binding motifs. These residues are not above reasonable contour in the high-resolution map, and we therefore did not include them in the model of the complex. (C, D and E) The three L protein-binding motifs of the phosphorprotein model are shown as sticks (carbon, magenta; nitrogen, blue; oxygen, red). The density map (EMD-20614, B sharpened map) is shown as gray mesh, and around the phosphorprotein only (carving radius = 2.0 Å). Note the absence of density for almost all acidic amino acid side chains (Asp and Glu, indicated by asterisks), which is characteristic of density maps obtained from cryo-EM reconstructions.

**Figure S5.**
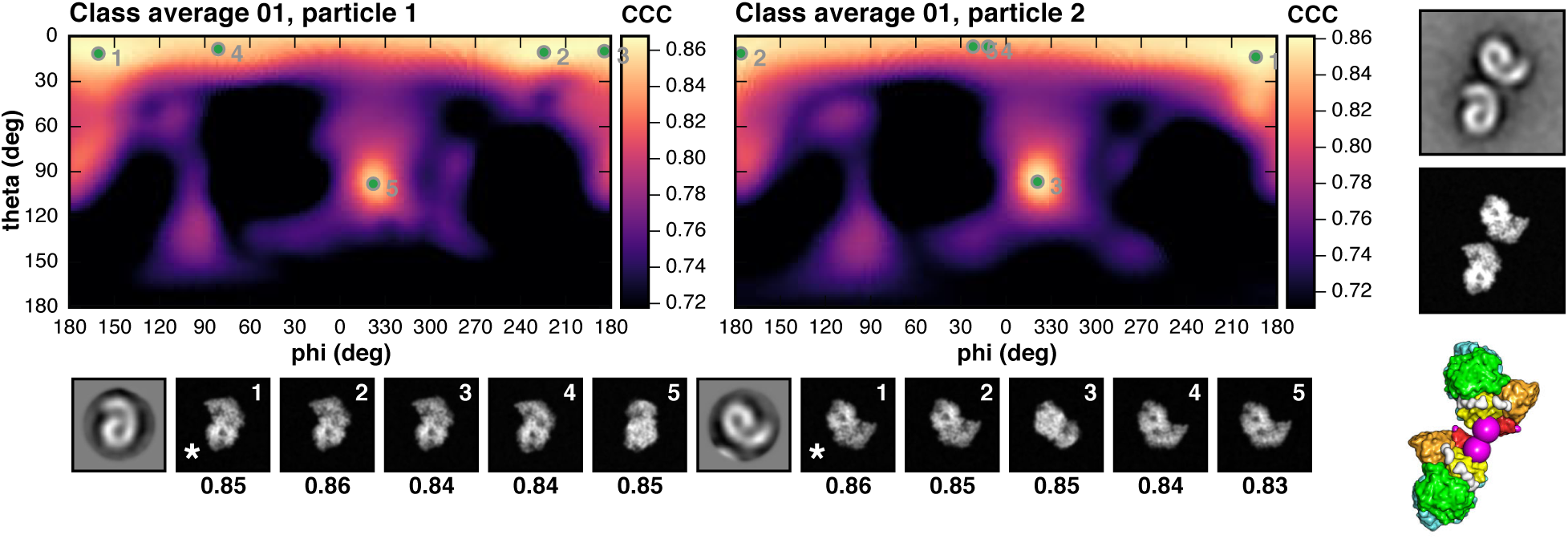
Projection Matching of Negative-Stain Class Averages. Related to Figure 3 and Data S3. The two heat maps show the cross-correlation coefficient (CCC) after aligning each protomer of the dimer to reference images projected when viewed from all possible angles (phi = 0–360°, theta = 0–180°). The top five peaks in each heat map are indicated by green dots and numbered; the corresponding references projections are shown at the bottom, together with the negative-stain particle image. The “correct” reference projection, as judged by eye, is labeled with and asterisk. On the right, the original class average (top), the calculated projection form two aligned references (middle), and the aligned model of the dimer (bottom) are shown. P protein residues 105 (the last residue observed in our structure) are shown as large magenta spheres. The figure shows the results for negative-stain class average 1 only. Data for all other classes can be found in Data S1.

**Figure S6.**
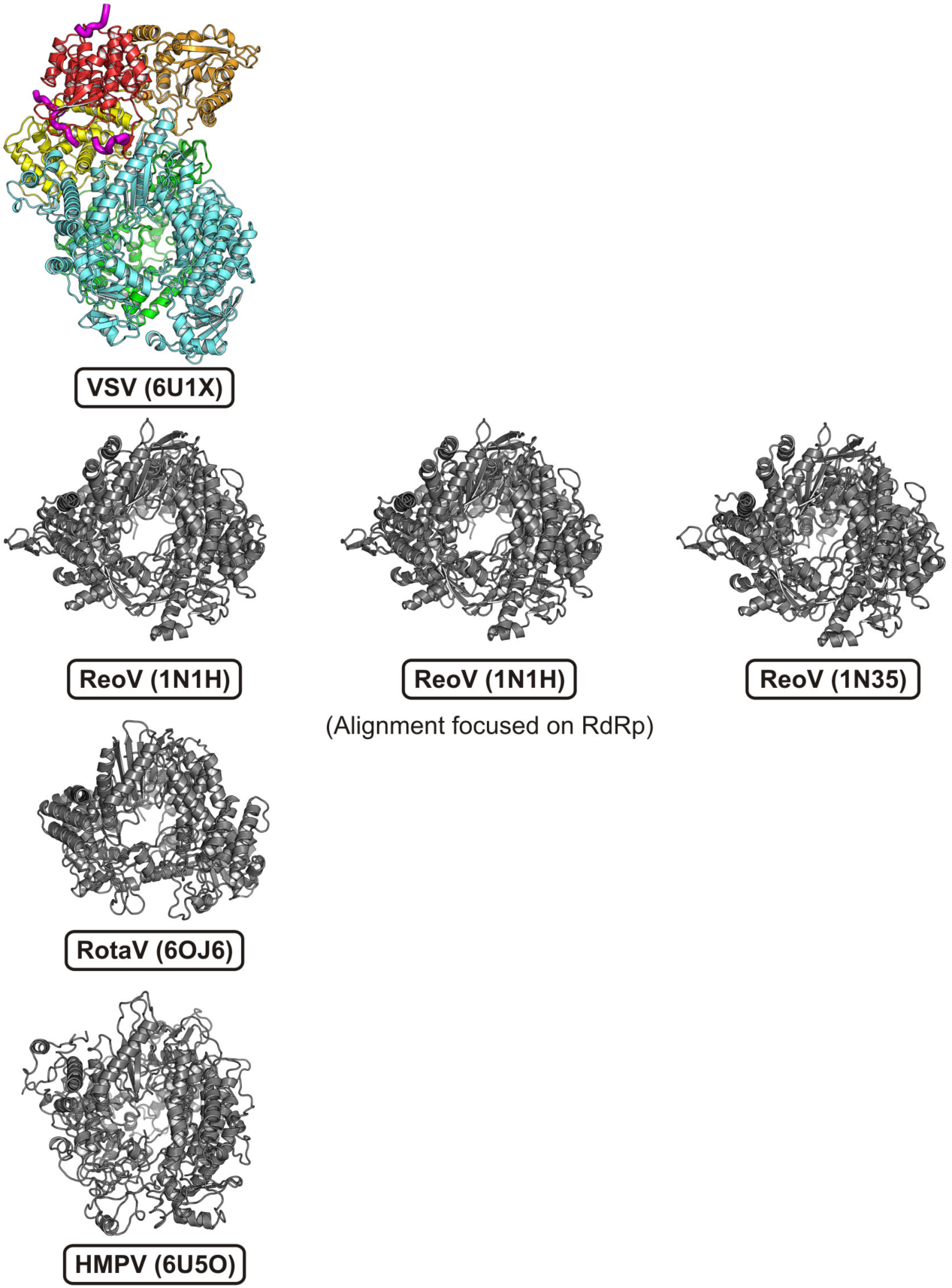
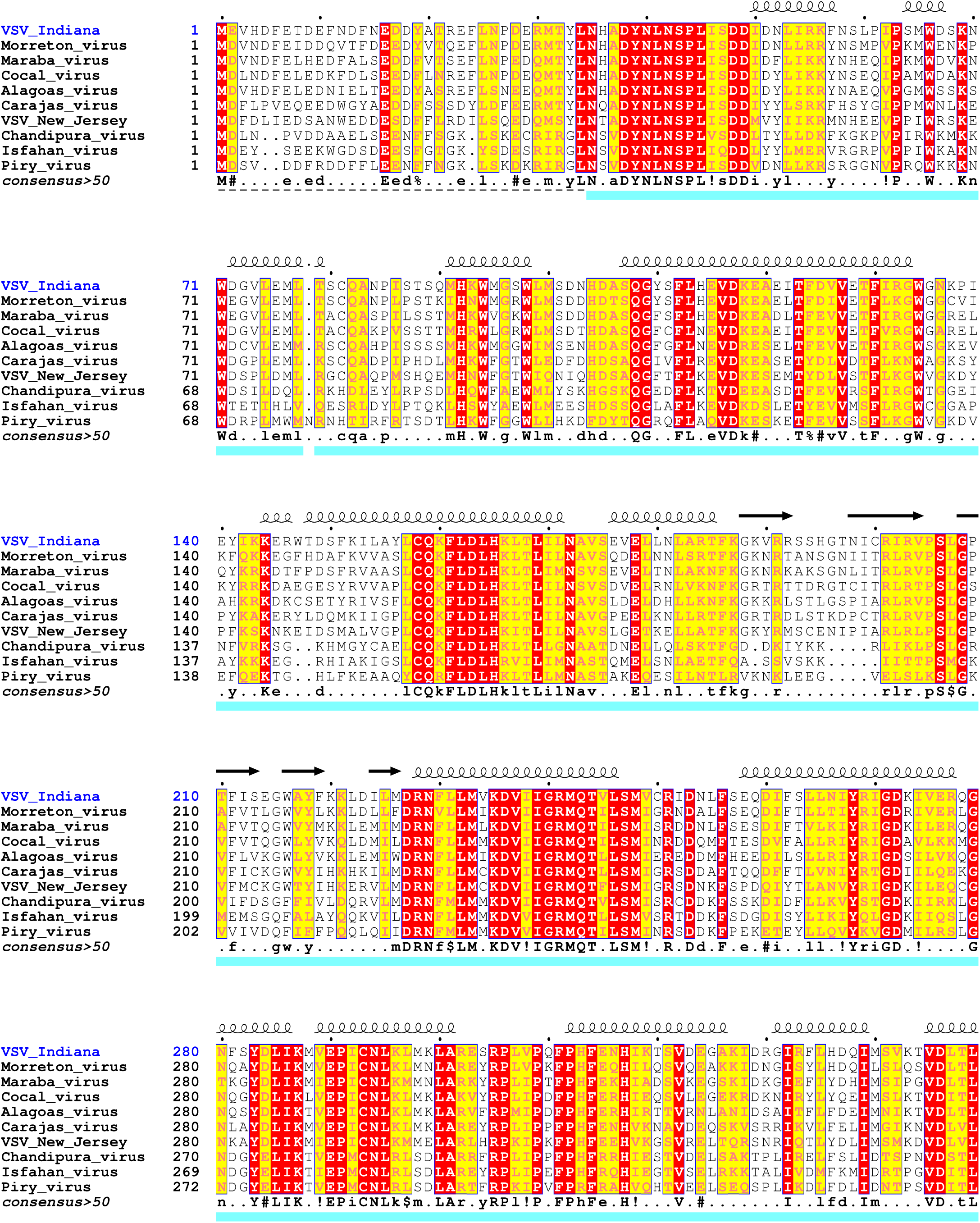
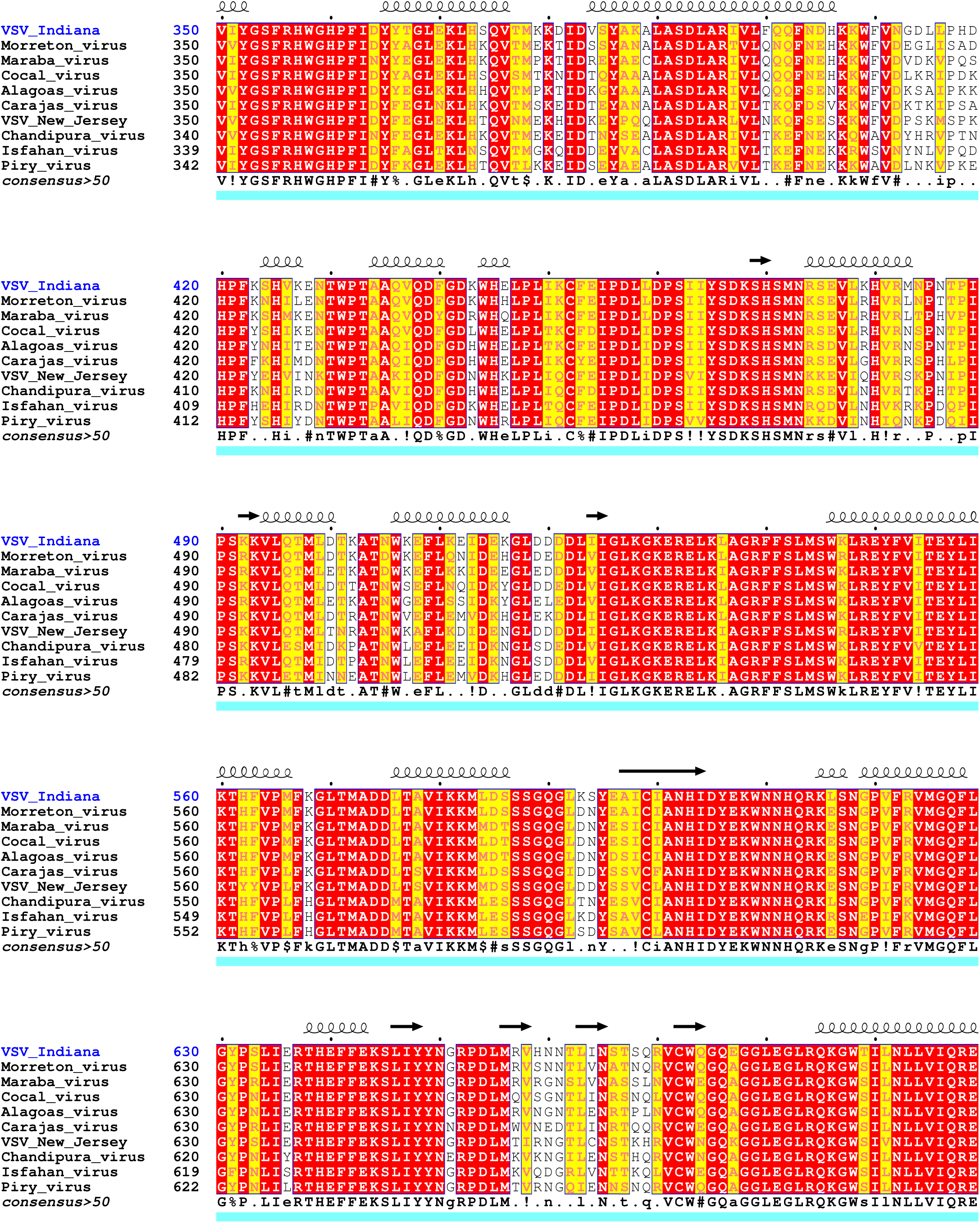
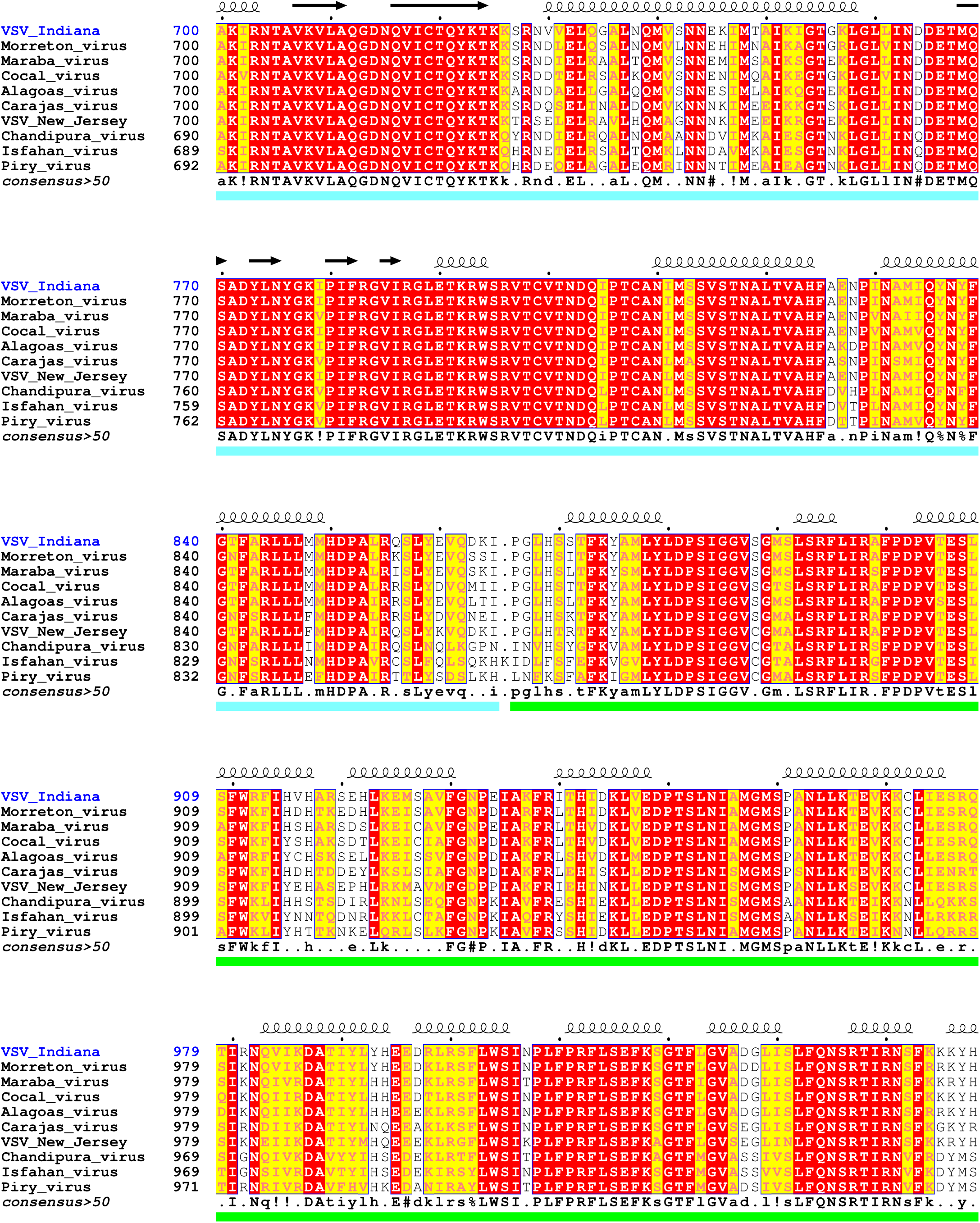
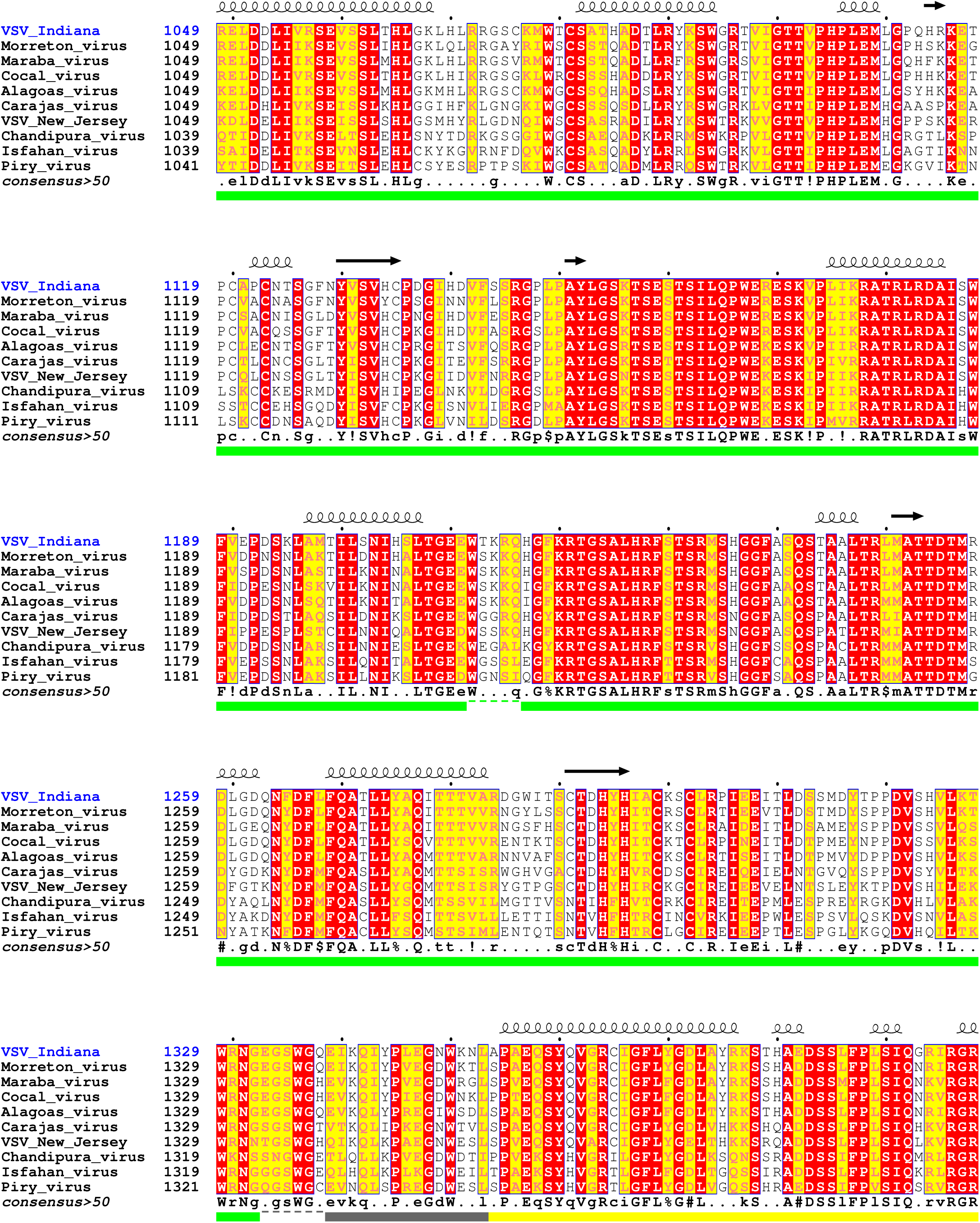
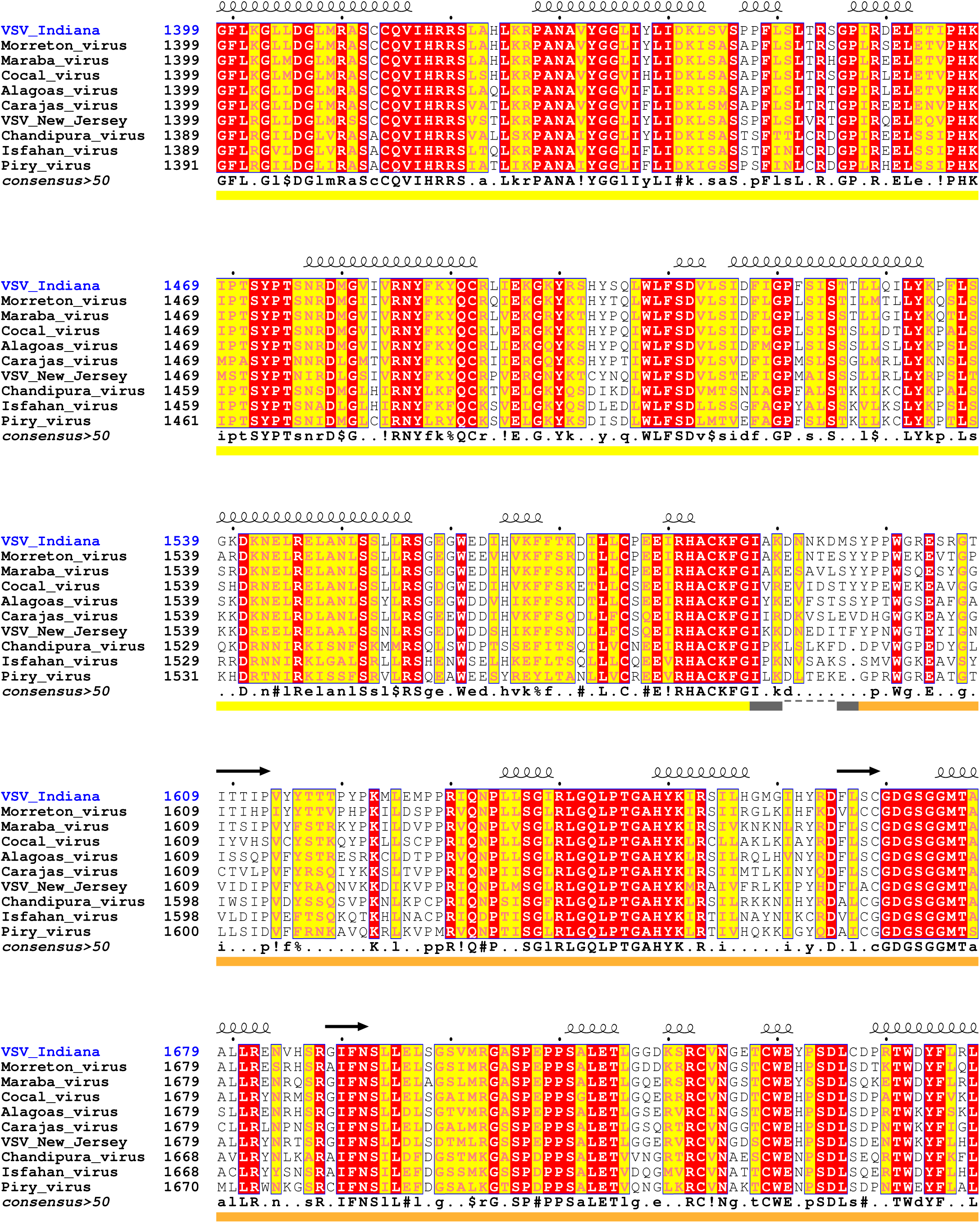
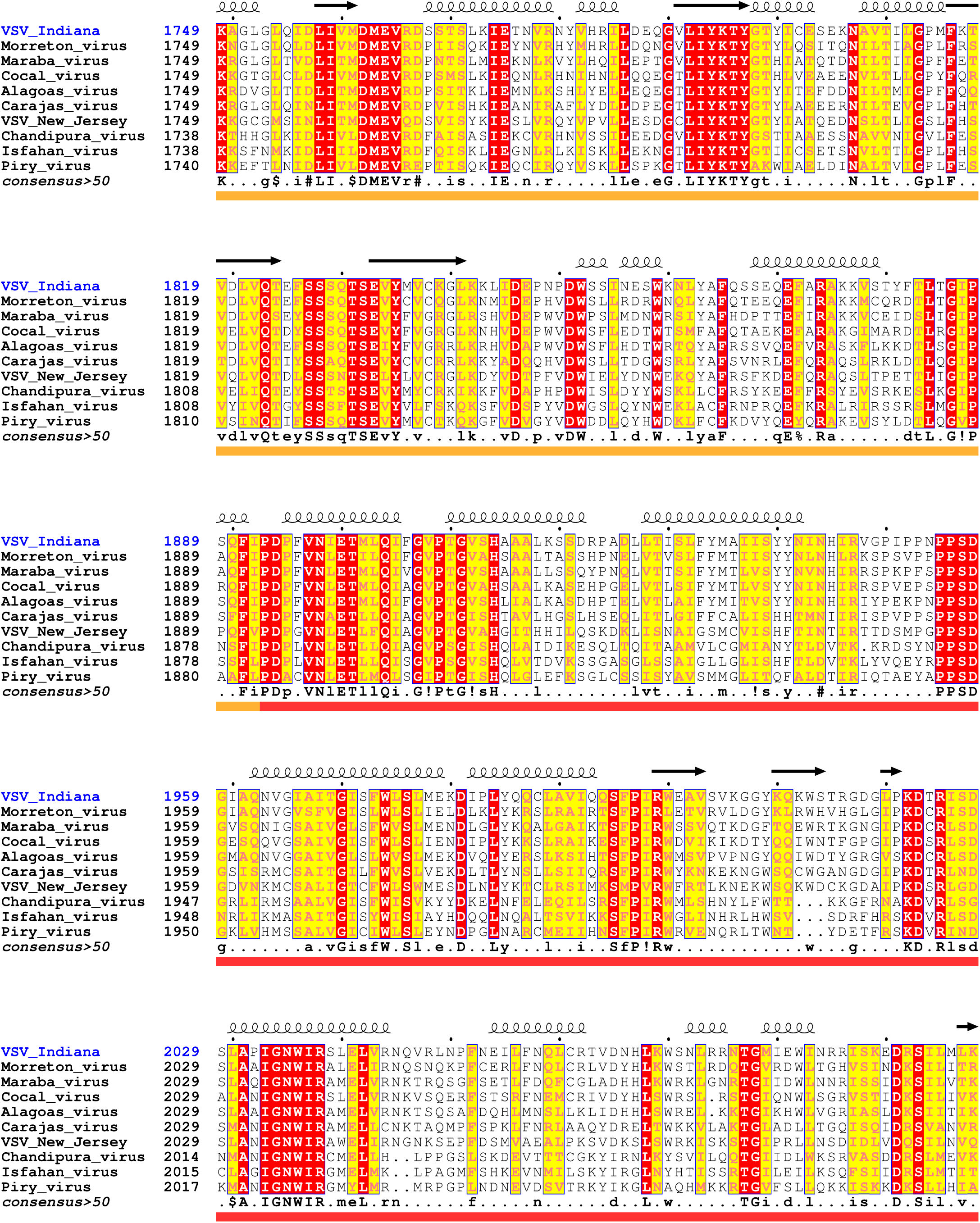
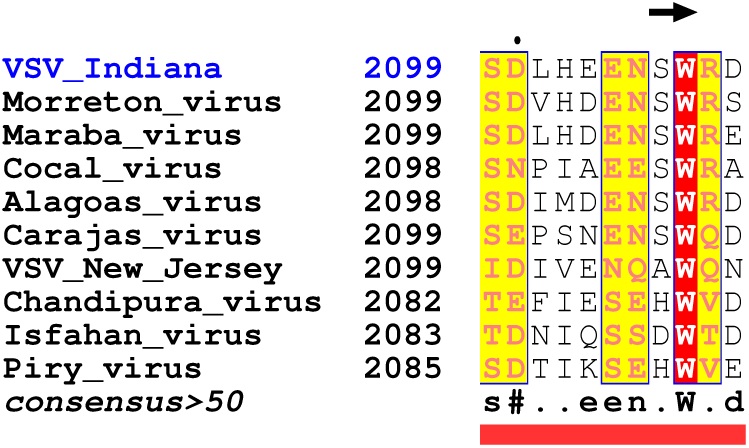
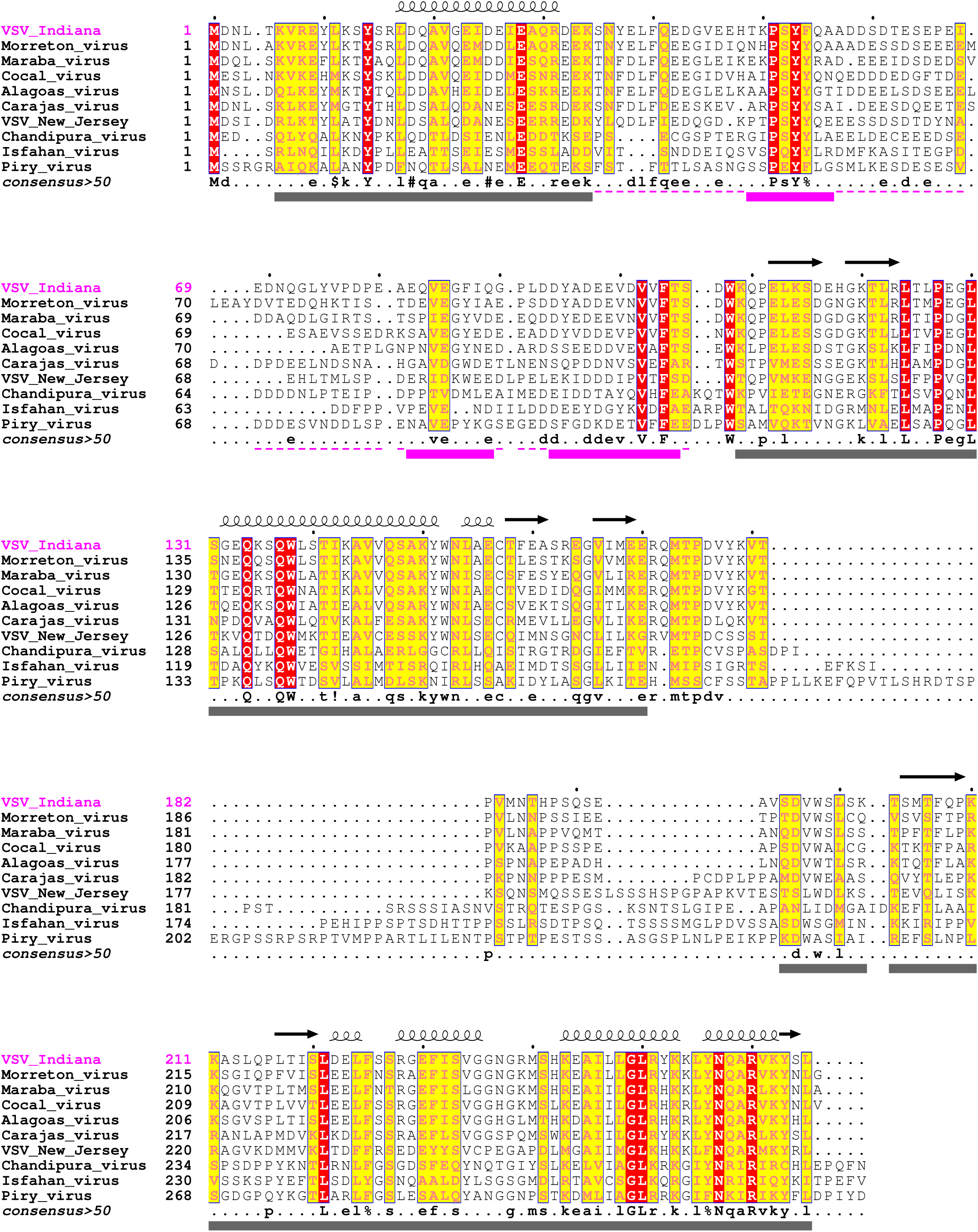
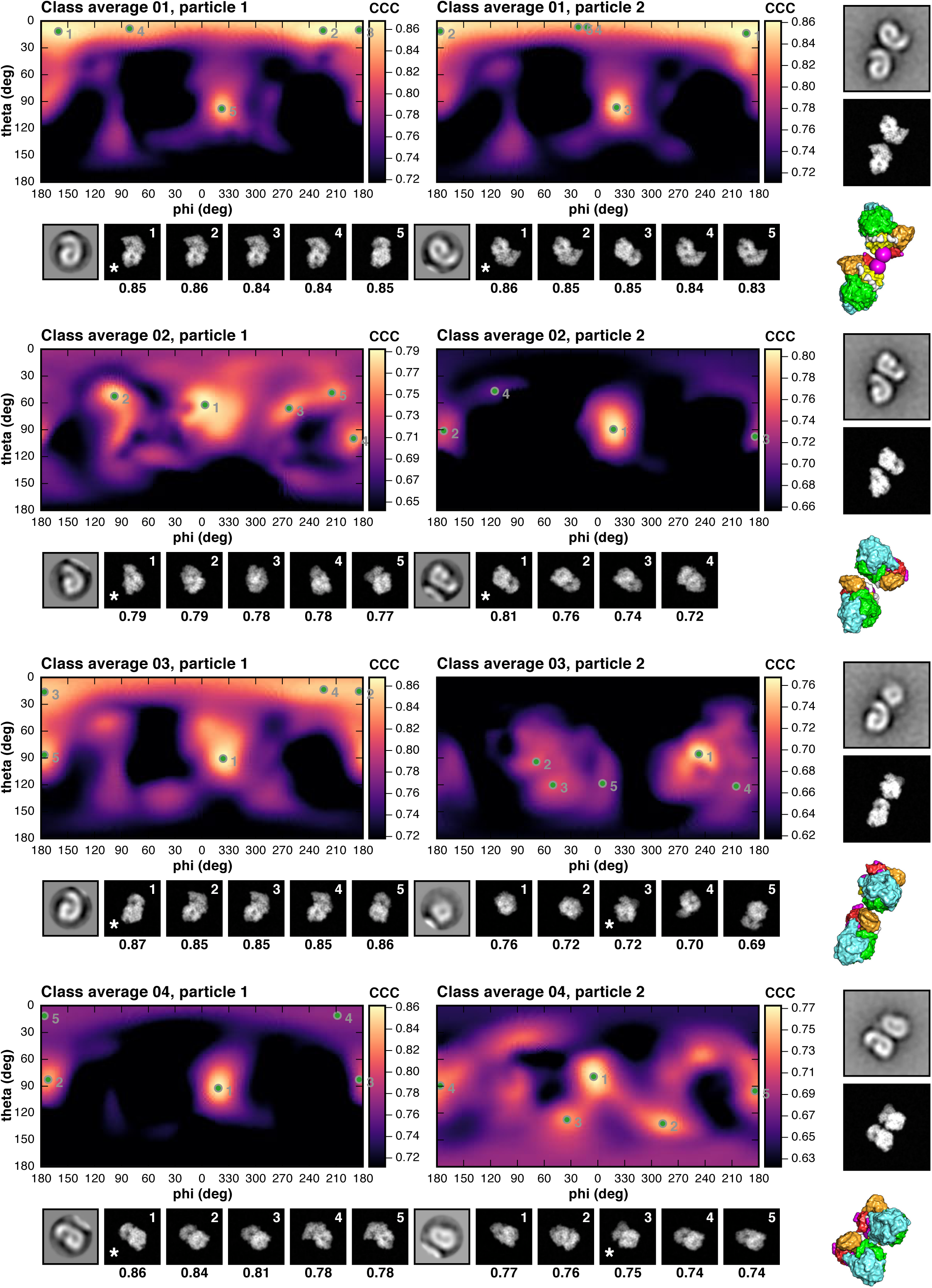
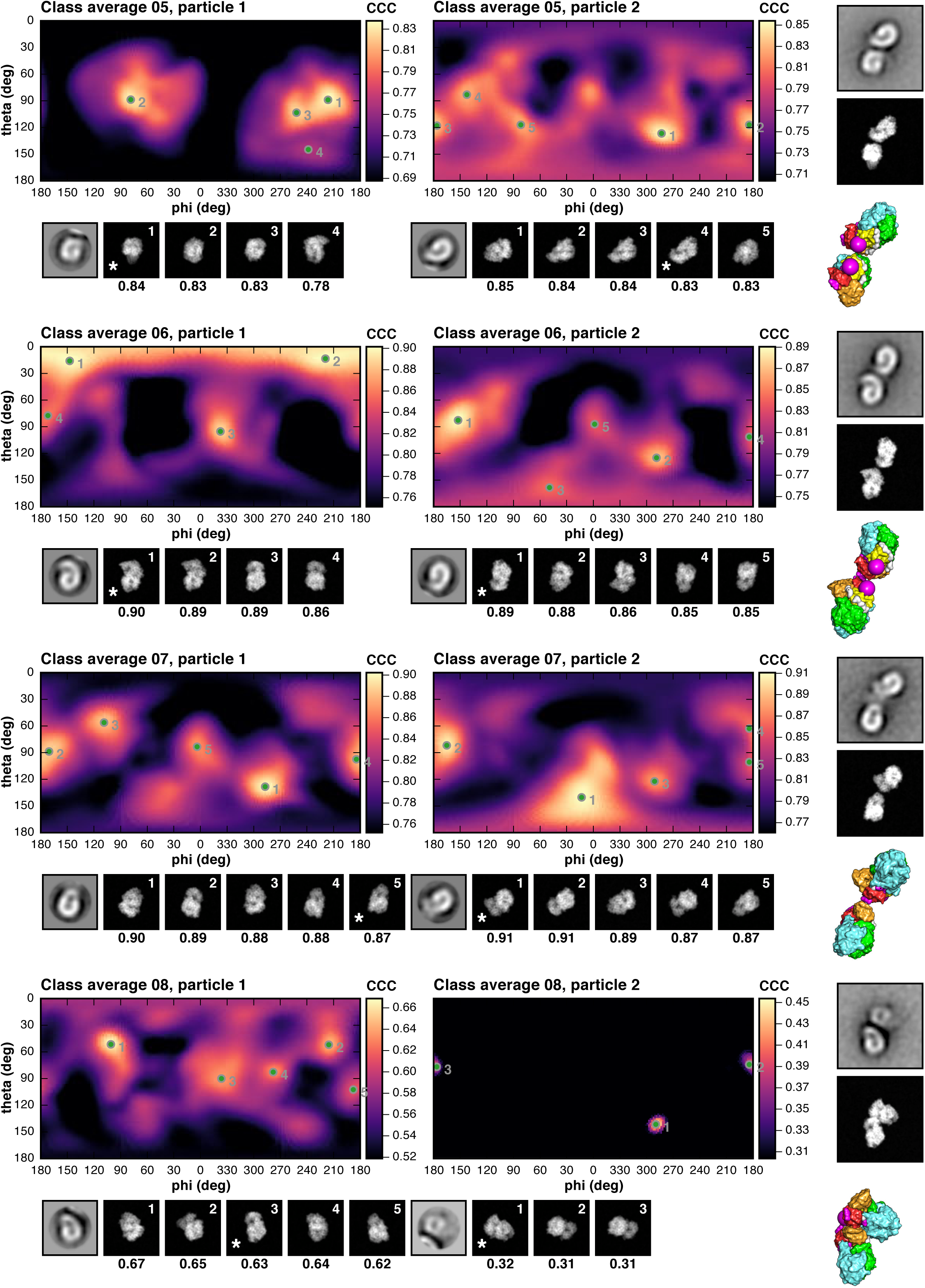
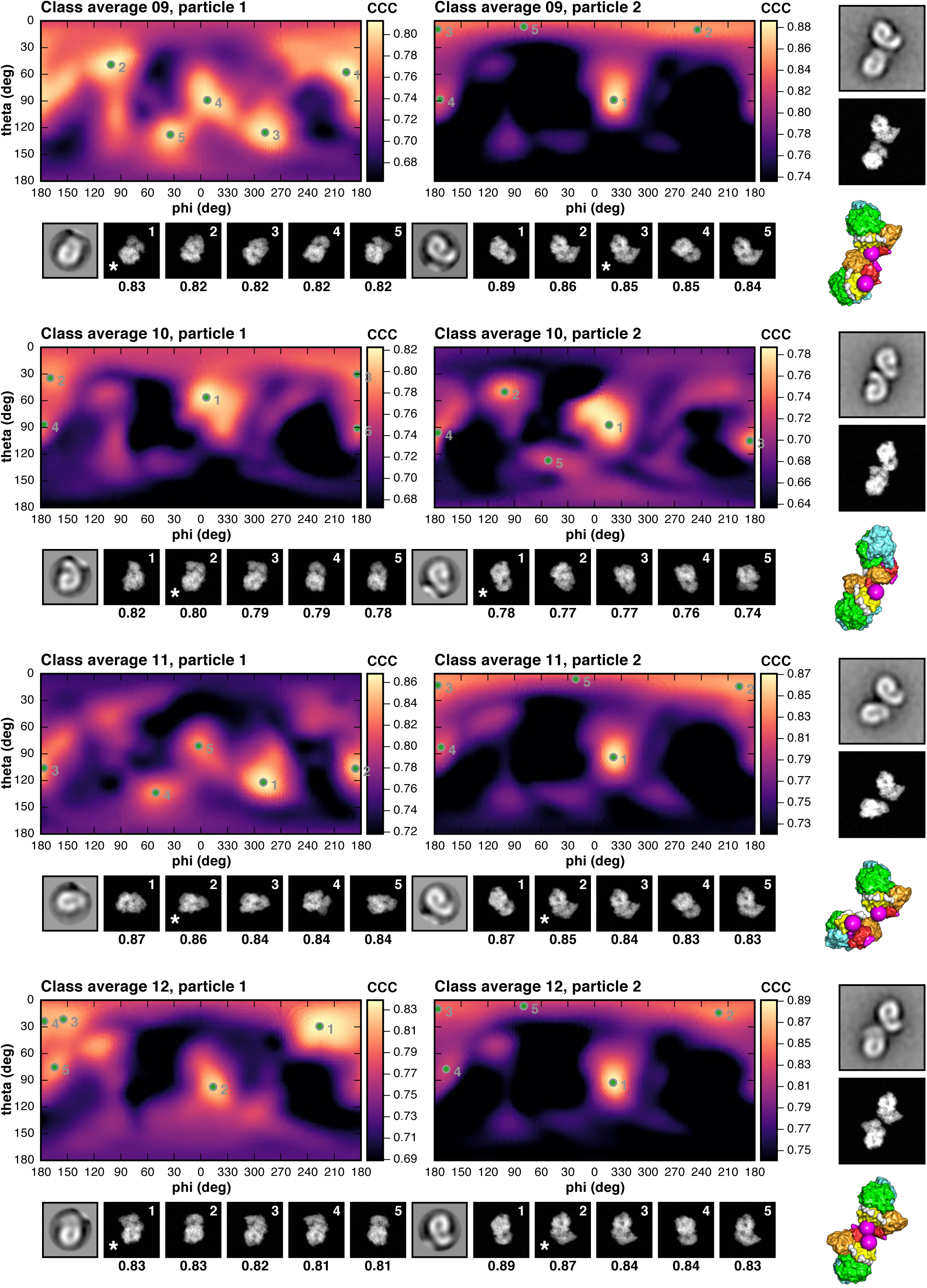
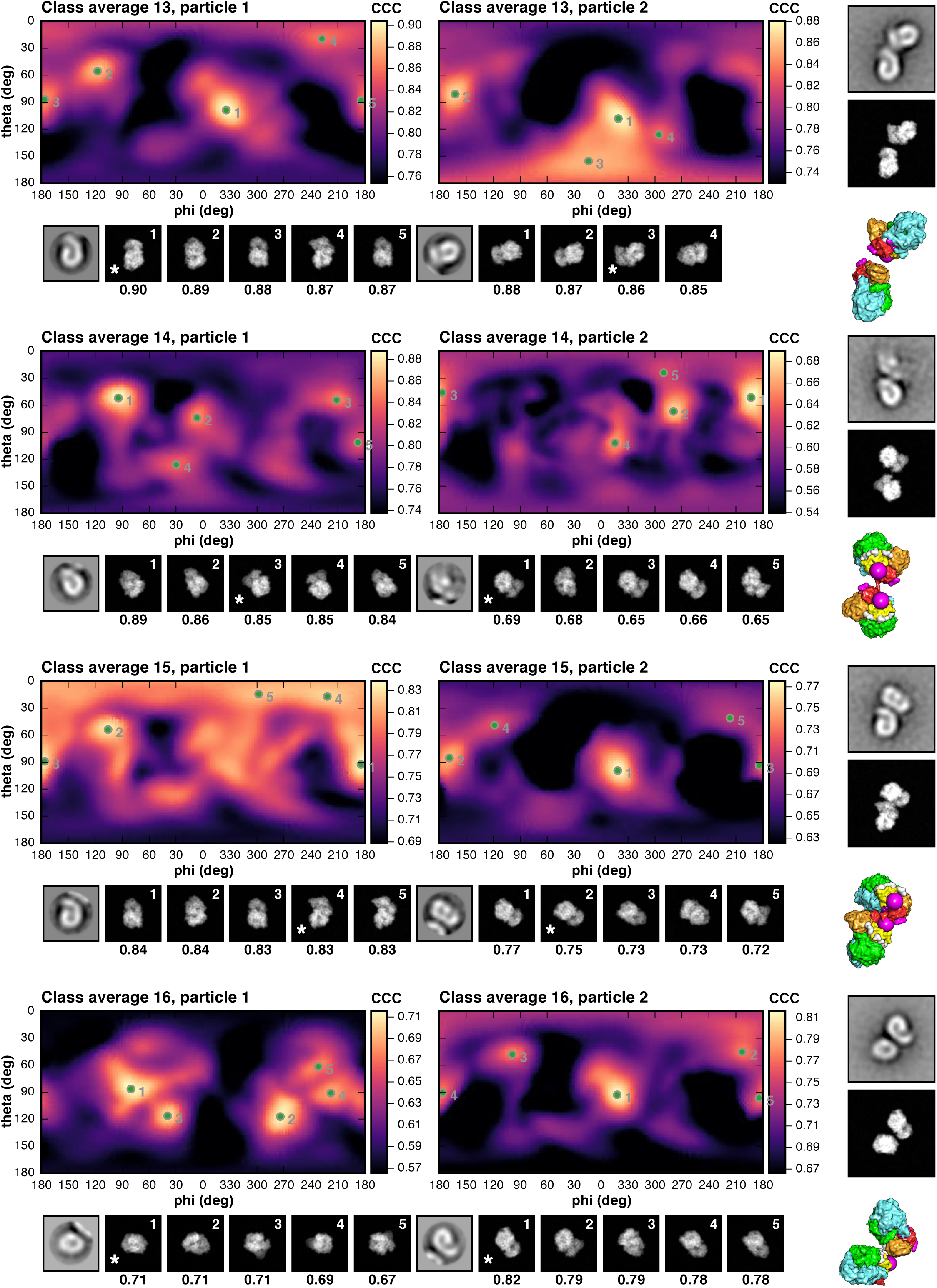
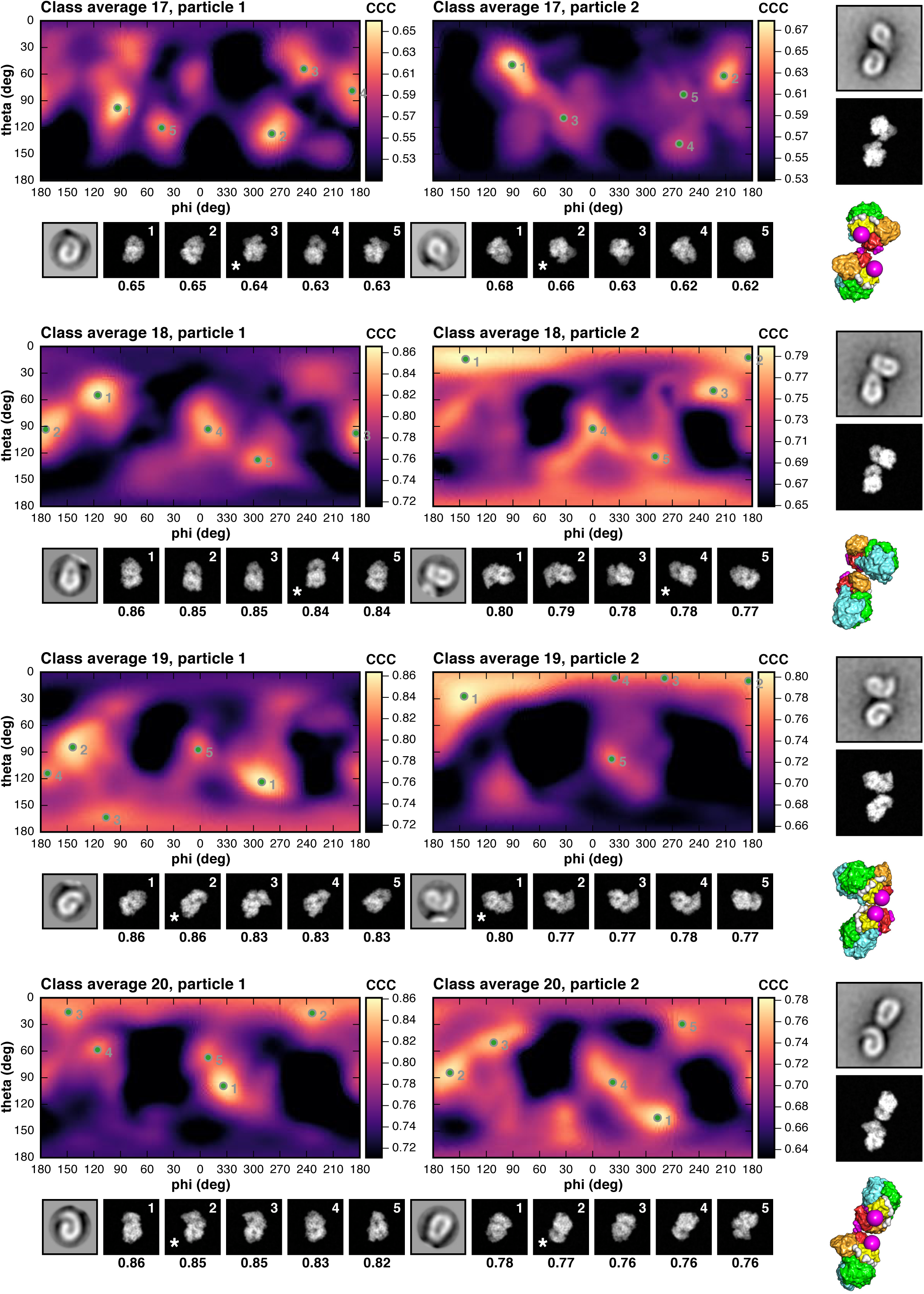
Superposition of Viral Polymerase Structures. Related to Figure 4, Table S3 and STAR Methods. Structures were superimposed as described in the Methods section. The polymerase structures are from vesicular stomatitis virus (VSV), reovirus (ReoV), rotavirus (RotaV), and human metapneumovirus (HMPV). Protein Data Bank (PDB) accession identifiers for the structures are given. See Table S3 for details of the structures and superpositions.

**Data S1. L Protein Multiple Sequence Alignment. Related to Figure 1.**

L protein amino acid sequences from different viruswere aligned with MAFFT (Katoh et al., 2002) and displayed with ESPript (Robert and Gouet, 2014). UniProt sequence accession identifiers are given in parenthesis: VSV_Indiana (P03523), vesicular stomatitis virus (Indiana strain); Morreton_virus (A0A0D3R1D1), Morreton vesiculovirus; Maraba_virus (F8SPF5), Maraba virus; Cocal_virus (B3FRL0), Cocal virus; Alagoas_virus (B3FRL5), vesicular stomatitis Alagoas virus; Carajas_virus (A0A0D3R1M8), Carajas virus; VSV_New_Jersey (P16379), vesicular stomatitis New Jersey virus; Isfahan_virus (P16379), Isfahan virus; Piry_virus (A0A1I9L1X2), Piry virus. Solid bars at the bottom of the sequences (domain-wise coloring as in Figure 1) are L protein residues modeled in our structure, dashed lines are residues included in our expression construct. Secondary structure elements assigned using DSSP (Kabsch and Sander, 1983) from our L protein structure are annotated in black on top of the sequences.

**Data S2. P Protein Multiple Sequence Alignment. Related to Figure 2.**

P protein amino acid sequences from different viruswere aligned with MAFFT (Katoh et al., 2002) and displayed with ESPript (Robert and Gouet, 2014). UniProt sequence accession identifiers are given in parenthesis: VSV_Indiana (Q8B0I3), vesicular stomatitis virus (Indiana strain); Morreton_virus (A0A0D3R1G8), Morreton vesiculovirus; Maraba_virus (F8SPF1), Maraba virus; Cocal_virus (B3FRK7), Cocal virus; Alagoas_virus (B3FRL2), vesicular stomatitis Alagoas virus; Carajas_virus (A0A0D3R1C6), Carajas virus; VSV_New_Jersey (A0A222ZEJ1), vesicular stomatitis New Jersey virus; Isfahan_virus (Q5K2K6), Isfahan virus; Piry_virus (Q01769), Piry virus. Magenta solid bars at the bottom of the sequences are P protein residues modeled in our structure, magenta dashed lines are residues included in our expression construct. Gray solid bars at the bottom of the sequences are P protein residues of domains for which structures have been determined in isolation: N-terminal domain (P_NTD_, PDB-ID 3PMK), oligomerization domain (P_OD_, PDB-ID 2FQM), and C-terminal domain (P_CDT_, PDB-ID 2K47). Secondary structure elements assigned using DSSP (Kabsch and Sander, 1983) from those structures are annotated in black on top of the sequences.

**Data S3. Projection Matching of Negative-Stain Class Averages. Related to Figure 3 and Figure S5.**

This file contains the analysis results of projection matching for all 20 negative-stain class averages. See STAR Methods and Figure S5 for a description of the procedure and figure panels.

**Table S1.** Cryo-EM Data Collection and Model Statistics. Related to STAR Methods.

| VSV L-P |  |
| --- | --- |
| EMD-20614<br>PDB-ID 6U1X |  |
| Indiana strain |  |
| **Data collection** |  |
| Electron microscope | Titan Krios |
| Magnification | 103092 |
| Voltage (kV) | 300 |
| Defocus range ($\mu\text{m}$)<sup>a</sup> | 0.5–2.0 |
| Pixel size ($\text{\AA}$) | 0.97 |
| **Reconstruction** |  |
| Number of images | 83979 |
| Box size (pixels) | 400 |
| Symmetry imposed | $C_1$ |
| Map resolution ($\text{\AA}$)<sup>b</sup> | 3.0 |
| Estimated $B$ factor ($\text{\AA}^2$)<sup>c</sup> | 75 |
| **Model statistics** |  |
| Refinement resolution ($\text{\AA}$) | 3.0 |
| CC (mask) | 0.82 |
| Model composition |  |
| Non-hydrogen atoms | 16761 |
| Protein residues | 2059 |
| Mean $B$ factor ($\text{\AA}^2$) | 70.7 |
| R.m.s. deviations |  |
| Bond lengths ($\text{\AA}$) | 0.004 |
| Bond angles ($^\circ$) | 0.548 |
| Validation |  |
| MolProbity clash score | 3.7 |
| Poor rotamers (%) | 0.1 |
| Ramachandran plot<sup>d</sup> |  |
| Favored (%) | 99.7 |
| Allowed (%) | 0.3 |
| Disallowed (%) | 0.0 |

<sup>a</sup> Approximate range of underfocus.
<sup>b</sup> Resolution where Fourier shell correlation (FSC) between half-maps drops below 0.143 after applying a loose spherical shell mask and scaling of the FSC to estimate what it would be for a tight mask (Grant, T., Rohou, A., and Grigorieff, N. (2018). *cisTEM*, user-friendly software for single-particle image processing. *Elife* 7).
<sup>c</sup> The map was sharpened using *cisTEM*, with simple flattening of the power spectrum. Applying a sharpening $B$ factor of $-75 \text{ \AA}^2$ to the unsharpened map would approximately reproduce the *cisTEM*-sharpened map.
<sup>d</sup> Ramachandran restraints were used in the refinement target.

**Table S2.** Improved Model Regions of PDB-ID 6U1X (3.0 Å Resolution) Compared to PDB-ID 5A22 (3.8 Å Resolution). Related to Figure 1 and STAR Methods.

| VSV L protein domain | Residue range <sup>a</sup> | Comment |
| --- | --- | --- |
| RdRp | 61–64 | Loop remodeled, sequence register shift of 60–64 |
|  | 414–430 | Sequence register shift of 414–430 |
|  | 587–593 | Loop remodeled, sequence register shift of 586–593 |
| Cap | 958 | Loop adjusted |
|  | 1035–1046 | Sequence register shift of 1035–1046 |
|  | 1072–1075 | Loop remodeled |
|  | 1100 | Loop adjusted |
|  | 1122–1125 | Loop remodeled, sequence register shift of 1124–1126 |
|  | 1164–1170 | Loop remodeled, sequence register shift of 1161–1170 |
|  | 1195–1232 | Sequence register shift of 1195–1227 |
|  | 1262 | Loop adjusted |
|  | 1314 | Loop adjusted |
|  | 1321–1332 | Region adjusted |
| CD | 1380–1395 | Region remodeled, sequence register shift of 1379–1396 |
|  | 1498 | Loop adjusted |
|  | 1535–1539 | Loop remodeled, sequence register shift of 1535–1538 |
| MT | 1718 | Loop adjusted |
|  | 1806–1851 | Sequence register shift of 1805–1841 |
| CTD | 1952–1955 | Loop remodeled, sequence register shift of 1950–1955 |
| | 2006–2008 | Shift in $\beta$ hairpin tip |
|  | 2066 | Loop remodeled |
|  | 2086–2094 | Loop remodeled, sequence register shift of 2084–2094 |
| | 2109 | Cα in main chain density instead side chain density of 2108 |

<sup>a</sup> Residue ranges with Cα distances larger than 3.0 Å between 6U1X and 5A22 are listed.

**Table S3.** Superposition Statistics of Viral Polymerase Structures. Related to Figure 4, Figure S6 and STAR Methods.

| Structure | PDB-ID | Program | Statistics |
| --- | --- | --- | --- |
| VSV L protein |  |  |  |
| Initiation-competent state | 6U1X | N.A. | N.A. |
| Reovirus λ3 polymerase | | | |
| Initiation complex | 1N1H | LSQMAN<sup>a</sup> | r.m.s. fit = 2.278 Å with 411 Cα atoms |
| | | O<sup>b</sup> | r.m.s. fit = 2.263 Å with 242 Cα atoms |
| Elongation complex | 1N35 | LSQMAN<sup>a</sup> | r.m.s. fit = 2.263 Å with 409 Cα atoms |
| Rotavirus VP1 |  |  |  |
| Elongation complex | 6OJ6 | LSQMAN<sup>a</sup> | r.m.s. fit = 2.134 Å with 381 Cα atoms |
| HMPV L protein |  |  |  |
| Initiation-competent state | 6U5O | LSQMAN<sup>a</sup> | r.m.s. fit = 2.015 Å with 858 Cα atoms |

<sup>a</sup> We obtained global alignments in LSQMAN with these commands:
``` fast m1 A m2 A 50 25 100 ! initial brute-force structural alignment on a coarse grid im m1 * m2 * ! intermediate-fit dp m1 a m2 a sq 3.5 10 ! final fit with Needleman–Wunsch algorithm ```
<sup>b</sup> We obtained the alignment in O with these commands:
``` lsq_exp ! initial superposition of active site residues (VSVL708–L719) lsq_imp ! improvement of the alignment by considering the whole structures ```

## REFERENCES

Afonine, P.V. (2017). phenix.mtriage: a tool for analysis and validation of cryo-EM 3D reconstructions. Computational Crystallography Newsletter 8, 25.

Afonine, P.V., Poon, B.K., Read, R.J., Sobolev, O.V., Terwilliger, T.C., Urzhumtsev, A., and Adams, P.D. (2018). Real-space refinement in PHENIX for cryo-EM and crystallography. Acta Crystallogr. D Struct. Biol. 74, 531–544.

Bell, J.M., Chen, M., Baldwin, P.R., and Ludtke, S.J. (2016). High resolution single particle refinement in EMAN2.1. Methods 100, 25–34.

Buchholz, U.J., Finke, S., and Conzelmann, K.K. (1999). Generation of bovine respiratory syncytial virus (BRSV) from cDNA: BRSV NS2 is not essential for virus replication in tissue culture, and the human RSV leader region acts as a functional BRSV genome promoter. J. Virol. 73, 251–259.

Chen, V.B., Arendall, W.B., 3rd, Headd, J.J., Keedy, D.A., Immormino, R.M., Kapral, G.J., Murray, L.W., Richardson, J.S., and Richardson, D.C. (2010). MolProbity: all-atom structure validation for macromolecular crystallography. Acta Crystallogr. D Biol. Crystallogr. 66, 12–21.

Ding, H., Green, T.J., Lu, S., and Luo, M. (2006). Crystal structure of the oligomerization domain of the phosphoprotein of vesicular stomatitis virus. J. Virol. 80, 2808–2814.

Fuerst, T.R., Niles, E.G., Studier, F.W., and Moss, B. (1986). Eukaryotic transient-expression system based on recombinant vaccinia virus that synthesizes bacteriophage T7 RNA polymerase. Proc. Natl. Acad. Sci. U. S. A. 83, 8122–8126.

Ge, P., Tsao, J., Schein, S., Green, T.J., Luo, M., and Zhou, Z.H. (2010). Cryo-EM model of the bullet-shaped vesicular stomatitis virus. Science 327, 689–693.

Gilman, M.S.A., Liu, C., Fung, A., Behera, I., Jordan, P., Rigaux, P., Ysebaert, N., Tcherniuk, S., Sourimant, J., Eleouet, J.F., et al. (2019). Structure of the Respiratory Syncytial Virus Polymerase Complex. Cell.

Górski, K.M., Hivon, E., Banday, A.J., Wandelt, B.D., Hansen, F.K., Reinecke, M., and Bartelmann, M. (2005). HEALPix: A Framework for High-Resolution Discretization and Fast Analysis of Data Distributed on the Sphere. Astrophys. J. 622, 759–771.

Grant, T., and Grigorieff, N. (2015). Measuring the optimal exposure for single particle cryo-EM using a 2.6 A reconstruction of rotavirus VP6. Elife 4, e06980.

Grant, T., Rohou, A., and Grigorieff, N. (2018). cisTEM, user-friendly software for single-particle image processing. Elife 7.

Green, T.J., and Luo, M. (2009). Structure of the vesicular stomatitis virus nucleocapsid in complex with the nucleocapsid-binding domain of the small polymerase cofactor, P. Proc. Natl. Acad. Sci. U. S. A. 106, 11713–11718.

Green, T.J., Zhang, X., Wertz, G.W., and Luo, M. (2006). Structure of the vesicular stomatitis virus nucleoprotein-RNA complex. Science 313, 357–360.

Hunter, J.D. (2007). Matplotlib: A 2D graphics environment. Comput. Sci. Eng. 9, 90–95.

Iverson, L.E., and Rose, J.K. (1981). Localized attenuation and discontinuous synthesis during vesicular stomatitis virus transcription. Cell 23, 477–484.

Jenni, S., Salgado, E.N., Herrmann, T., Li, Z., Grant, T., Grigorieff, N., Trapani, S., Estrozi, L.F., and Harrison, S.C. (2019). In situ Structure of Rotavirus VP1 RNA-Dependent RNA Polymerase. J. Mol. Biol.

Jones, T.A., Zou, J.Y., Cowan, S.W., and Kjeldgaard, M. (1991). Improved methods for building protein models in electron density maps and the location of errors in these models. Acta Crystallogr. A 47, 110–119.

Kabsch, W., and Sander, C. (1983). Dictionary of protein secondary structure: pattern recognition of hydrogen-bonded and geometrical features. Biopolymers 22, 2577–2637.

Katoh, K., Misawa, K., Kuma, K., and Miyata, T. (2002). MAFFT: a novel method for rapid multiple sequence alignment based on fast Fourier transform. Nucleic Acids Res. 30, 3059–3066.

Kleywegt, G.J., and Jones, T.A. (1994). Detection, delineation, measurement and display of cavities in macromolecular structures. Acta Crystallogr. D Biol. Crystallogr. 50, 178–185.

Kleywegt, G.J., and Jones, T.A. (1997). Detecting folding motifs and similarities in protein structures. Methods Enzymol. 277, 525–545.

Kranzusch, P.J., and Whelan, S.P. (2011). Arenavirus Z protein controls viral RNA synthesis by locking a polymerase-promoter complex. Proc. Natl. Acad. Sci. U. S. A. 108, 19743–19748.

Li, J., Rahmeh, A., Morelli, M., and Whelan, S.P. (2008). A conserved motif in region v of the large polymerase proteins of nonsegmented negative-sense RNA viruses that is essential for mRNA capping. J. Virol. 82, 775–784.

Liang, B., Li, Z., Jenni, S., Rahmeh, A.A., Morin, B.M., Grant, T., Grigorieff, N., Harrison, S.C., and Whelan, S.P.J. (2015). Structure of the L Protein of Vesicular Stomatitis Virus from Electron Cryomicroscopy. Cell 162, 314–327.

Lu, X., McDonald, S.M., Tortorici, M.A., Tao, Y.J., Vasquez-Del Carpio, R., Nibert, M.L., Patton, J.T., and Harrison, S.C. (2008). Mechanism for coordinated RNA packaging and genome replication by rotavirus polymerase VP1. Structure 16, 1678–1688.

Mastronarde, D.N. (2005). Automated electron microscope tomography using robust prediction of specimen movements. J. Struct. Biol. 152, 36–51.

Morin, B., Rahmeh, A.A., and Whelan, S.P. (2012). Mechanism of RNA synthesis initiation by the vesicular stomatitis virus polymerase. EMBO J. 31, 1320–1329.

Ogino, M., Gupta, N., Green, T.J., and Ogino, T. (2019). A dual-functional priming-capping loop of rhabdoviral RNA polymerases directs terminal de novo initiation and capping intermediate formation. Nucleic Acids Res. 47, 299–309.

Ongradi, J., Cunningham, C., and Szilagyi, J.F. (1985). The role of polypeptides L and NS in the transcription process of vesicular stomatitis virus New Jersey using the temperature-sensitive mutant tsE1. J. Gen. Virol. 66 (Pt 5), 1011–1023.

Pattnaik, A.K., and Wertz, G.W. (1990). Replication and amplification of defective interfering particle RNAs of vesicular stomatitis virus in cells expressing viral proteins from vectors containing cloned cDNAs. J. Virol. 64, 2948–2957.

Rahmeh, A.A., Morin, B., Schenk, A.D., Liang, B., Heinrich, B.S., Brusic, V., Walz, T., and Whelan, S.P. (2012). Critical phosphoprotein elements that regulate polymerase architecture and function in vesicular stomatitis virus. Proc. Natl. Acad. Sci. U. S. A. 109, 14628–14633.

Rahmeh, A.A., Schenk, A.D., Danek, E.I., Kranzusch, P.J., Liang, B., Walz, T., and Whelan, S.P. (2010). Molecular architecture of the vesicular stomatitis virus RNA polymerase. Proc. Natl. Acad. Sci. U. S. A. 107, 20075–20080.

Robert, X., and Gouet, P. (2014). Deciphering key features in protein structures with the new ENDscript server. Nucleic Acids Res. 42, W320–324.

Rohou, A., and Grigorieff, N. (2015). CTFFIND4: Fast and accurate defocus estimation from electron micrographs. J. Struct. Biol. 192, 216–221.

Shuman, S. (1997). A proposed mechanism of mRNA synthesis and capping by vesicular stomatitis virus. Virology 227, 1–6.

Speir, J.A., and Johnson, J.E. (2012). Nucleic acid packaging in viruses. Curr. Opin. Struct. Biol. 22, 65–71.

Tao, Y., Farsetta, D.L., Nibert, M.L., and Harrison, S.C. (2002). RNA synthesis in a cage–structural studies of reovirus polymerase lambda3. Cell 111, 733–745.

Tekes, G., Rahmeh, A.A., and Whelan, S.P. (2011). A freeze frame view of vesicular stomatitis virus transcription defines a minimal length of RNA for 5’ processing. PLoS Pathog 7, e1002073.

van Heel, M., Harauz, G., Orlova, E.V., Schmidt, R., and Schatz, M. (1996). A new generation of the IMAGIC image processing system. J. Struct. Biol. 116, 17–24.

Wertz, G.W., Whelan, S., LeGrone, A., and Ball, L.A. (1994). Extent of terminal complementarity modulates the balance between transcription and replication of vesicular stomatitis virus RNA. Proc. Natl. Acad. Sci. U. S. A. 91, 8587–8591.

Whelan, S.P., Ball, L.A., Barr, J.N., and Wertz, G.T. (1995). Efficient recovery of infectious vesicular stomatitis virus entirely from cDNA clones. Proc. Natl. Acad. Sci. U. S. A. 92, 8388–8392.

Whelan, S.P., Barr, J.N., and Wertz, G.W. (2000). Identification of a minimal size requirement for termination of vesicular stomatitis virus mRNA: implications for the mechanism of transcription. J. Virol. 74, 8268–8276.

Winn, M.D., Ballard, C.C., Cowtan, K.D., Dodson, E.J., Emsley, P., Evans, P.R., Keegan, R.M., Krissinel, E.B., Leslie, A.G., McCoy, A., et al. (2011). Overview of the CCP4 suite and current developments. Acta Crystallogr. D Biol. Crystallogr. 67, 235–242.

Zivanov, J., Nakane, T., Forsberg, B.O., Kimanius, D., Hagen, W.J., Lindahl, E., and Scheres, S.H. (2018). New tools for automated high-resolution cryo-EM structure determination in RELION-3. Elife 7.

